# Modulation of gluconic acid metabolism enhances nanofibrillated bacterial cellulose production from agro-industrial by-products

**DOI:** 10.1101/2025.08.25.672120

**Authors:** Ryo Takahama, Go Takayama, Miho Suginaka, Yuma Ishido, Shunsuke Nagai, Kazuki Nagai, Masato Uenishi, Kenji Tajima

**Author notes:** Corresponding author: Kenji Tajima. Author contributions: R. T.: Research plan, reactor experiments on the WT strain, culture data analysis, and manuscript writing; G. T.: NMR and WAXD data analyses, and AFM image analysis; M. S.: Gene knockout experiments; Y. I.: Reactor experiments on GM strains and culture data analysis; S. N.: Gene knockout experiments; K. N.: Gene knockout experiments; M. U.: Reactor preliminary experiments on the WT strain; K. T.: Supervision. All authors have read and approved the final manuscript.

## Abstract

Efficient and robust bacterial cellulose production is essential for advancing the sustainable bioeconomy. In this study, we investigated the impacts of metabolism of organic acids, mainly gluconic acid (GA), on nanofibrillated bacterial cellulose (NFBC) production by *Komagataeibacter intermedius* NEDO-01 under various culture conditions in aerated stirred-tank reactors. In cultures of the wild-type strain in a standard medium, rapid GA production decreased the medium pH and depleted glucose, inhibiting cell growth and reducing the NFBC yield. However, proper pH control and continuous feeding reversed these effects, resulting in a 3-fold increase in NFBC yield (from 2.45 to 7.59 g/L). In cultures of a glucose dehydrogenase gene-deficient (Δ*gcd*) strain, lack of a pH drop and glucose depletion facilitated better cell growth, yielding 1.85-times more NFBC than that in wild-type cultures under pH-uncontrolled no-feed conditions (4.53 g/L). Notably, GA supplementation accelerated cell growth but significantly inhibited NFBC synthesis, suggesting that GA uptake redirects the carbon flux toward central metabolism.

In the corn steep liquor (Csl)-based medium, cell growth was significantly enhanced, and NFBC yield was equivalent to or higher than that obtained with the Hestrin–Shramm medium. GA accumulation was markedly reduced, suppressing pH fluctuation. Under these optimized conditions, three molasses types were tested with Csl, yielding relatively high NFBC. Structural analysis of NFBC produced using these alternative media revealed slight differences in the fiber width distribution, with crystallinity and fiber width remaining constant. Overall, NFBC of consistent quality can be produced in stirred-tank reactors using *Komagataeibacter* spp. from various agricultural by-products.

**Importance:** In this study, we investigated the interplay between organic acid metabolism and nanofibrillated bacterial cellulose (NFBC) production in stirred-tank reactor (STR) cultures of *Komagataeibacter intermedius* NEDO-01. While it is well known that gluconic acid production competes with cellulose biosynthesis in *Komagataeibacter*, the quantitative relationship between these pathways under varying culture conditions has not been fully elucidated. By applying optimized feeding strategies and employing a glucose dehydrogenase knockout mutant, we demonstrated that suppressing gluconic acid accumulation significantly enhances NFBC yield. Furthermore, we explored the use of agro-industrial by-products, including molasses and corn steep liquor, as alternative, low-cost feedstocks. Structural characterization confirmed that NFBCs produced under these conditions maintained consistent quality. These findings contribute to the development of scalable, cost-effective microbial production processes for nanocellulose, which is essential for advancing the sustainable bioeconomy.

**Key Points:**

- GA accumulation inhibited growth and cellulose production by *Komagataeibacter*
- Enhanced central metabolism elevated NFBC yield but reduced its production rate
- Consistent NFBC properties were achieved in STRs using various by-product sources

## Introduction

Nanofibrillated bacterial cellulose (NFBC) is a type of surface-modified cellulose nanofibril produced directly from *Komagataeibacter* cultures in aerated stirred-tank reactors (STRs). It consists of two cellulosic components: Native bacterial cellulose (BC) and cellulose derivatives, such as hydroxypropyl (HPC) and carboxymethyl (CMC) cellulose. When cellulose-producing bacteria, such as *Komagataeibacter intermedius* NEDO-01 (NEDO-01), are cultured in the presence of cellulose derivatives, such as HPC and CMC, these additives modify the surface of newly synthesized BC microfibrils, resulting in the formation of NFBC in a highly dispersed state (Kose et al. 2013; Tajima et al. 2017). Polymers, such as HPC, are stably adsorbed onto the NFBC surface, allowing surface chemical modifications using these surface polymers as scaffolds for chemical reactions in common organic solvents (Hashim et al. 2022, 2024). NFBC is characterized by a long fiber length and high aspect ratio (≥1000) (Tsujisaki et al. 2024), which confer shear-thinning properties that protect cells in animal cell cultures (Kaneko et al. 2024).

Unlike plant-derived cellulose nanofibrils, NFBC can be produced from sugars using conventional STRs, facilitating its continuous scalable production. Furthermore, its surface chemical properties can be tuned by selecting different cellulose derivatives, offering significant advantages for material design and functionalization. However, various unused resources, including crude and low-purity feedstocks, must be used for its cost-effective production. Whether NFBC can be consistently produced from such resources remains unclear.

*K. intermedius* NEDO-01 (NEDO-01) acts as a production strain due to its high cellulose-producing capacity, metabolic stability, and strong tolerance to impurities commonly found in industrial and agricultural by-products (Kose et al. 2013). Kose *et al*. revealed that NEDO-01 produces NFBC using waste glycerol, a biodiesel fuel by-product containing lipids, fatty acids, and methanol, as a carbon source. In contrast, NFBC productivities of other representative cellulose-producing strains, such as ATCC53582 and ATCC23769, are significantly low under the same conditions (Kose et al. 2013). These characteristics make NEDO-01 a promising strain for industrial-scale NFBC production.

In wild-type *Komagataeibacter* cultures, by-products such as organic acids and water-soluble polysaccharides (acetan) are produced in addition to cellulose. Notably, large amount of gluconic acid (GA) is rapidly generated in the presence of glucose as the carbon source (Rezazadeh 2020). This phenomenon, in which sugars and alcohols (e.g., ethanol) are rapidly oxidized into organic acids, leading to a sharp drop in pH, is known as oxidative fermentation, which is characteristic process of the *Acetobacteraceae* family. Although oxidative fermentation is energetically inefficient and inhibits the growth of producing organisms, it acts as a competitive survival strategy in nutrient-rich environments (Matsushita 2004). Suppressing the formation of such by-products, including GA and acetan, increases cellulose production in *Komagataeibacter* cultures (Kuo et al. 2015; Ryngajłło et al. 2019; Montenegro-Silva et al. 2024).

In this study, we examined the effects of various culture conditions on the production of NFBC and organic acids, particularly GA, during aerated STR cultivation. We further evaluated the roles of GA production and utilization in NFBC production using a glucose dehydrogenase (*gcd*) knockout mutant. Under optimized conditions, we investigated the effects of three types of molasses of variable origin and purity on the NFBC yield, structure, and metabolic behavior of NEDO-01. The resulting NFBC types were characterized via solid-state nuclear magnetic resonance (NMR) spectroscopy, wide-angle X-ray diffraction (WAXD), and atomic force microscopy (AFM) to assess the effects of the culture conditions on the molecular structure, crystallinity, and nanofiber morphology of NFBC.

## Materials and methods

### Materials

*K. intermedius* NEDO-01 (NITE P-1495), which was isolated in our previous work (Kose et al. 2013), was used as the production strain. HPC (grade [M]; Mn: 620,000; molecular substitution [MS]: 3.6; NISSO, Tokyo, Japan) was selected as the dispersing agent. Chemicals for culture media preparation were purchased from Kanto Chemical (Tokyo, Japan; D-glucose, D-fructose, citric acid, and Na_2_HPO_4_) and Thermo Fischer Scientific (Waltham, MA, USA; Bacto yeast extract [Gibco] and Bacto peptone [Gibco]), with purity ≥ 98%. Three types of molasses (sugar beet molasses grades A [molasses A] and B [molasses B], and sugar cane molasses [molasses SC]) were kindly provided by Nippon Beet Sugar Manufacturing Co., Ltd. (Tokyo, Japan). Corn steep liquor (Csl) was purchased from Kogo Starch Co., Ltd. (Chiba, Japan). HCl, H2SO4, and NaOH were purchased from Kanto Chemical with purity ≥ 98%. 5-Fluorocytosine was purchased from Tokyo Chemical Industry (Tokyo, Japan). Cellulase ONOZUKA R-10 was purchased from Yakult Pharmaceutical Industry CO., Ltd. (Tokyo, Japan).

### Pre-treatment of molasses and Csl

Molasses A, B, and SC (all provided as 80°Bx solutions) were diluted to 60°Bx with deionized water. Molasses A was mixed with a 1/50 volume of 2.4 M HCl and heated in an 80 °C water bath for 45 min. Molasses B and SC were mixed with 96% H_2_SO_4_ until reaching pH 2.0 and heated in an 80 °C water bath for 3 and 5 h, respectively. After treatment, resulting liquid samples were collected and diluted 10 times with deionized water, and sugar compositions were analyzed via high-performance liquid chromatography. Typically, hydrolysate of molasses A contains 300_350 mg/mL glucose, whereas those of molasses B and SC contain approximately 200 mg/mL glucose. All hydrolyzed molasses were stored at 4 °C and used within one month.

Csl (pH ∼4.0) was mixed with 5 M NaOH until reaching pH 6.5, heated in a 70 °C water bath for 3 h, and centrifuged at 11,200 × *g* for 15 min at 25 °C to remove the suspended solids. The supernatant was collected and stored at 4 °C for less than one month until use.

### Culture medium composition

A culture medium was prepared as follows based on the Hestrin–Shramm (HS) medium composition, with some modifications: 12.5 g/L glucose, 12.5 g/L fructose, 5 g/L yeast extract, 5 g/L peptone, 2.7 g/L Na_2_HPO_4_, and 1.5 g/L citric acid. The feeding solution was prepared as a 62.5 g/L glucose and 62.5 g/L fructose solution in water. When molasses was used as the main carbon source, glucose and fructose were replaced with molasses containing equivalent concentrations of glucose (fructose concentrations were slightly lower than glucose concentrations in molasses). When Csl was used as the nitrogen source, 5 g/L peptone was replaced with 50 g/L Csl. The pH of the culture medium was adjusted using a 2 M NaOH or 0.5 M H_2_SO_4_ aqueous solution. The final compositions are listed in Table 1.

**Table 1.** Amounts of carbon and nitrogen sources in different culture media (YE, yeast extract; Csl, corn steep liquor).

| Abbreviation | Carbon sources |  |  | Nitrogen sources |  |  |  |  |
| --- | --- | --- | --- | --- | --- | --- | --- | --- |
|  | Glucose<br>(g L<sup>-1</sup>) | Fructose<br>(g L<sup>-1</sup>) | Molasses<br>(mL L<sup>-1</sup>) | YE<br>(g L<sup>-1</sup>) | Peptone<br>(g L<sup>-1</sup>) | Csl<br>(g L<sup>-1</sup>) | Na<sub>2</sub>HPO<sub>4</sub><br>(g L<sup>-1</sup>) | Citric acid<br>(g L<sup>-1</sup>) |
| HS(GF) | 12.5 | 12.5 | - | 5 | 5 | - | 2.7 | 1.5 |
| Csl(GF) | 12.5 | 12.5 | - | 5 | - | 50 | 2.7 | 1.5 |
| Csl(A) | - | - | ~180 | - | - | 50 | 2.7 | 1.5 |
| Csl(B) | - | - | ~300 | - | - | 50 | 2.7 | 1.5 |
| Csl(SC) | - | - | ~340 | - | - | 50 | 2.7 | 1.5 |

### Construction of a *gcd* gene-deficient (**Δ***gcd*) mutant via markerless gene deletion

Markerless deletion method with two-step homologous recombination was used to delete *gcd* from the NEDO-01 genomic DNA (gDNA). Gene knockout target *gcd* was selected based on its amino acid homology with that reported in previous studies (Montenegro-Silva et al. 2024). Gene deletion plasmid was constructed based on pUC19 and designed to incorporate the blue pigment indigoidine synthase-related genes, *bpsA* and *pptA*, for the visual selection of recombinant colonies based on the dark blue color (Takahashi et al. 2007). A counterselection system using *codA* and *codB*, which are used in acetic acid bacteria (Kostner et al. 2017; Bimmer et al. 2022; Bimmer et al. 2023), was also incorporated into our system. Homology arms were designed to be homologous to approximately 1,200-bp region upstream and downstream of the target gene. Plasmids and primers used in this study are listed in **Tables S1** and **S2** respectively. The plasmid design is shown in **Fig. S1** and the construction method is described in *Supplementary material* (supplementary method). Plasmids constructed in this study will be deposited to Addgene and made publicly available upon publication (Addgene ID: 246236).

The constructed plasmids were introduced into the electro cells (Warren 2011) of NEDO-01 via electroporation using a 0.1-cm cuvette (Bio-Rad, Hercules, CA, USA) at 18 kV/cm field strength (Hall 1992), spread onto the HS agar medium containing 100 μg/mL of ampicillin (Amp) with 200 μL of 3.5 M (NH_4_)_2_SO_3_, and incubated at 30 °C for three days. Dark-blue colonies were selected, suspended in the HS liquid medium, and spread onto the HS agar medium containing 100 μg/mL of Amp with 200 μL of 3.5 M (NH_4_)_2_SO_3_ for recombinant purification. The obtained colonies were inoculated into 10 mL of HS liquid medium containing 100 μg/mL Amp and 0.3% (w/v) cellulase and incubated at 30 °C and 150 rpm for three days. Then, gDNA was extracted from the bacteria. Plasmid insertion into the target region gDNA (first recombination) was confirmed via polymerase chain reaction (PCR).

After first recombination, the colonies were inoculated into 10 mL of HS liquid medium containing 200 µg/mL Amp and 0.3% (w/v) cellulase, incubated at 30 °C and 150 rpm for six days with shaking, spread onto the HS agar medium containing 240 µg/mL of fluorocytosine (FC) (Bimmer et al. 2022), and incubated at 30 °C for four days. The resulting white colonies were inoculated into 10 mL of HS liquid medium containing 240 µg/mL of FC and 0.3% (w/v) cellulase and grown at 30 °C and 150 rpm for three days. These cultures were further seeded onto the HS agar medium containing 240 µg/mL of FC, and the recombinant strain was carefully separated. Selected colonies were inoculated into 10 mL of HS liquid medium, incubated at 30 °C and 150 rpm for three days, and gDNA was extracted. Target gene deletion was confirmed via PCR using gDNA as a template, followed by PCR product sequencing with appropriate primers.

### Inoculum preparation

Frozen stock of *K. intermedius* NEDO-01 was thawed in a 25 °C water bath. Stock culture (250 μL) was inoculated into 10 mL of fresh HS medium supplemented with cellulase ONOZUKA R-10 (0.3% [w/v]) in a 50-mL flask with baffles (seed culture). This seed culture was incubated under rotary shaking (150 rpm) at 30 °C for 72 h, inoculated to 250 mL of fresh HS medium containing 1.5% (w/v) of HPC in a 1-L flask with baffles (IWAKI AGC TECHNO GLASS CO., Ltd, Shizuoka, Japan) at an inoculation rate of 5% (v/v), and cultivated under shaking at 150 rpm and 30 °C for 72 h (pre-culture).

### Culture method

Next, the pre-culture (250 mL) was inoculated into 5 L of fresh medium containing 2% (w/v) HPC in a 10-L bioreactor (ABLE Corporation, Tokyo, Japan). Temperature and airflow were set at 30 °C and 5 L/min (1.0 vvm), respectively. Atmospheric pressure inside the tank was set as slightly positive (+ 0.02 MPa). Dissolved oxygen was controlled at a target of 30% of saturation by maintaining the agitation speed within 300–500 rpm. Initial pH was adjusted to 5.2 by adding 2 M NaOH and 1 M H_2_SO_4_ aqueous solutions during the selective culture period (typically after 24 h). Approximately 15 mL of the culture sample was collected every 12 h during culture.

### Sugar concentration and optical density (OD) measurements

Each culture sample was diluted 10-times with Milli-Q water, and OD was measured at 600 nm (UV-1900i, Shimadzu, Kyoto, Japan). Each diluted culture sample was centrifuged at 3,000 × *g* for 5 min at 25 °C, and the supernatant was filtered via ultrafiltration. Sugar content was measured via size-exclusion chromatography using the COSMOSIL Sugar-D column (4.6ID × 250 mm; Nacalai Tesque, Inc., Kyoto, Japan) and a refractive index detector.

### NFBC purification

Each culture sample (10 g) was transferred to a 50-mL polypropylene centrifuge tube. Milli-Q water was added to each tube up to 30 mL, thoroughly mixed, and centrifuged at 18,000 × *g* for 30 min at 25 °C. The supernatant was discarded, and the precipitate containing NFBC was redispersed in 15 mL of Milli-Q water. An equal volume (15 mL) of 4% (w/v) NaOH aqueous solution was added to each culture tube. The tubes were shaken at 150 rpm and 70 °C for 2 h to lyse the bacteria and denature the proteins. After cooling, the sample was neutralized by adding a 3 M H_2_SO_4_ aqueous solution and centrifuged at 18,000 × *g* for 30 min at 25 °C to collect NFBC. The supernatant was discarded, and the precipitated NFBC pellet was washed and resuspended in ultrapure water (Milli-Q) and centrifuged. Finally, the supernatant was discarded, and the sample was dried at 105 °C for 2 h. Dry weight of each sample was measured after cooling.

### Culture data analysis

Cell dry weight (CDW) was estimated from the OD_600_ values using two conversion factors depending on the culture medium, as cell morphology differed between the HS-based- and Csl-based media. A factor of 0.216 mg/mL per OD_600_ unit was used for the HS-based medium, whereas a factor of 0.291 mg/mL per OD_600_ unit was used for the Csl-based medium. These values were derived from calibration curves based on the parallel measurements of OD_600_ and gravimetrically determined biomass. To determine the biomass weight, the culture samples were treated with 0.3% (w/v) cellulase ONOZUKA R-10 at 50 °C for 1 h to digest NFBC. The contribution of NFBC to OD_600_ was considered negligible and thus ignored.

Specific NFBC production and growth rates were calculated at 12-h intervals based on the CDW changes estimated from the OD_600_ values and NFBC concentrations. Specific growth rate was calculated using the following equation:

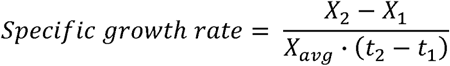

where *X_1_* and *X_2_* are the CDWs (g/L) at times *t_1_* and *t_2_*, respectively, and *X_avg_* is the average biomass during this time interval.

Specific NFBC production rate was calculated using the following equation:

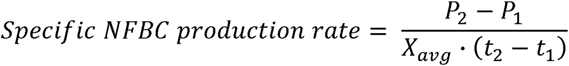

where *P_1_* and *P_2_* are the NFBC concentrations (g/L) at times *t_1_* and *t_2_*, respectively.

### AFM characterization of NFBC

Purified NFBC sample was diluted to 0.003% (w/v) with ultrapure water, gently sonicated (40 kHz) for 30 min, and placed on a mica template. NFBC was observed using AFM5000II (Hitachi High-Tech Corporation, Tokyo, Japan) equipped with the SI-DF3P2 microcantilever (Hitachi) in the dynamic mode. Height image was captured at a field of view of 20 μm × 20 μm with a resolution of 4096 × 4096 pixels. AFM images were analyzed using the Gwyddion (Nečas and Klapetek 2012) analysis software, and line profiles of the cross sections were obtained to evaluate the fiber width and height. In total, 160 fibers were measured for each sample. AFM measurements are subject to the tip convolution effect, which is the blurring of the image due to the tip shape of the probe. The tip convolution effect was corrected by setting the tip size to 10 nm and half angle to 17.5°, as previously indicated (Canet-Ferrer et al. 2014).

### X-ray diffraction

WAXD measurements were performed to evaluate the crystal structures of the prepared NFBC types. NFBC samples were freeze-dried, pressed into pellets, and subjected to X-ray diffraction measurements using Smart lab (Rigaku, Tokyo, Japan). All measurements were performed in the reflection mode using CuKa1 (wavelength = 1.54 Å) as the X-ray source, and scan range, scan speed, and scan resolution were set at 5–40°, 2°/min, and 0.01, respectively. The measured data were baseline-corrected with straight lines passing through two points at 5° and 30°. Peak fitting was also performed on the (100), (010), and (110) planes of the cellulose Iα crystal, and peak positions and full-width at half-maximum for these planes were analyzed. Peak fitting was performed using Python (version 3.12), and each peak was fitted using the Voigt function. Plane spacing was estimated from the peak positions, and crystallite size of each plane was estimated using the Scherrer equation.

### Solid-state NMR spectroscopy

Solid-state cross-polarization magic-angle spinning (MAS) ^13^C NMR measurements were performed to characterize the NFBC sample structures. Freeze-dried NFBC samples were packed into a 3.2-mm ZrO_2_ MAS rotor (Bruker, Billerica, MA, USA), and solid-state ^13^C NMR spectra were recorded at 125 MHz for ^13^C using the Avance Neo 500 spectrometer (Bruker).

MAS rate was set to 10 kHz, and duration of the 90° pulse for ^1^H, contact time for cross-polarization, and delay time between repetitions were set to 4.2 μs, 2 ms, and 4 s, respectively. Glycine was used as an external standard to calibrate the chemical shift scale relative to tetramethylsilane, and glycine carbonyl peak was assigned a chemical shift of 176.5 ppm. The crystallinity index (C4 crystal/[C4 crystal + C4 amorphous]) was calculated from the areas of the crystalline and amorphous C4 peaks in the obtained spectra. Cellulose Iα fraction was determined by separating the C1 peak into cellulose Iα and Iβ.

## Results

### Effect of GA accumulation on NFBC production by the wild-type strain

Oxidative fermentation is a characteristic feature of acetic acid bacterial metabolism, in which sugars or alcohols in the culture medium are rapidly oxidized to produce organic acids. This process involves several periplasmic dehydrogenases (Matsushita et al. 2004). Glucose is rapidly converted into GA in the *Komagataeibacter* genus (Rezazadeh 2020).

As shown in Fig. 1a, GA accumulation reached the maximum level (10 g/L) after 24 h in the *K. intermedius* NEDO-01 culture. Specific NFBC production rate reached its highest value at 48 h (**Fig. 1b**). Most glucose in the culture medium was consumed quickly within the initial 24 h, followed by fructose consumption (**Fig. 1c**). Cells continued to grow during the culture period (**Fig. S2a**), but pH and respiratory quotient were decreased after GA accumulation (**Fig. S2b** and **c**), indicating that the low pH due to GA accumulation reduced the growth and metabolic activity of *K. intermedius*. GA was gradually consumed as cultivation continued, and the cells resumed growth, with pH remaining low (3.6) due to citric acid accumulation (**Figs. 1a** and **S2b**). GA was entirely consumed in seven days, and the resulting NFBC yield was 2.45 g/L (**Fig. 1a**).

**Fig. 1.**
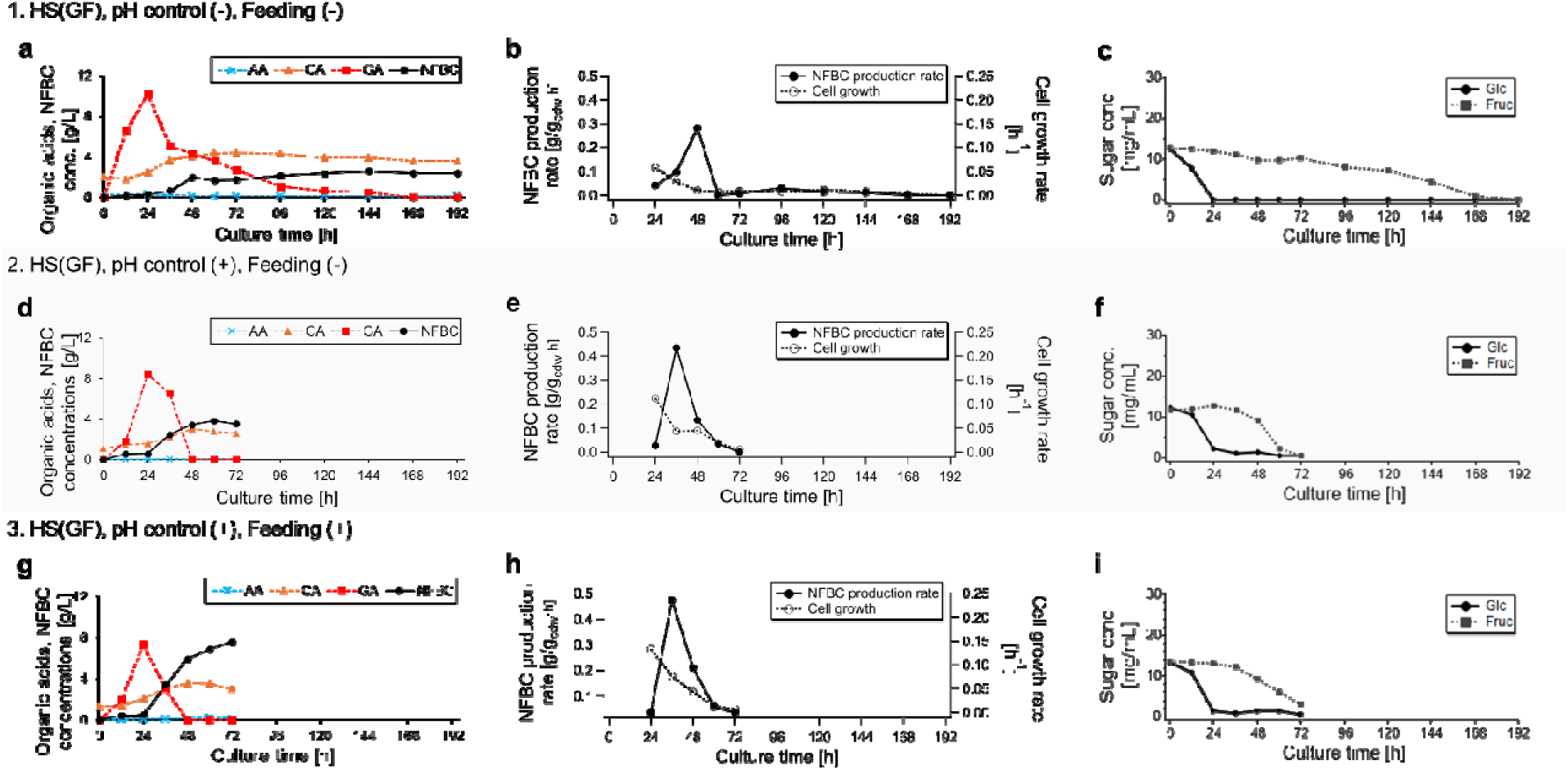
Time courses of nanofibrillated bacterial cellulose (NFBC) and organic acid accumulation (**a**, **d**, and **g**), specific NFBC production and growth rates (**b**, **e**, and **h**), and sugar concentrations (**c**, **f**, and **i**) in culture media. All experiments used the wild-type *K. intermedius* NEDO-01 strain and Hestrin–Shramm (HS[GF]) medium. Culture conditions were as follows: Normal batch culture without pH control or feeding (condition 1; **a**–**c**); pH regulation (24–72 h) using 2 M NaOHaq and 0.5 M H_2_SO_4_ (condition 2; **d–f**); a combination of pH control (24–72 h) and feeding (24–60 h) at a rate of 7.0 mL/h (condition 3; **g–i**). Specific NFBC production and growth rates were estimated from the optical density at 600 nm (OD_600_) values, and data points at 12 h were omitted due to low OD values

Adjusting the pH to the initial level (5.2) after 24 h resulted in a high NFBC yield (3.5 g/L) over a short culture period (72 h; **Fig. 1d**, condition 2). The maximum value of specific NFBC production rate (0.44 g/g_cdw_ h at 36h; **Fig. 1e**) was 1.6-fold higher than that in the normal batch culture (0.28 g/g_cdw_ h at 48h; **Fig. 1b**). GA and sugar consumption were also promoted (**Fig. 1d** and f). Other related data are shown in Fig. S2d-f.

Constant feeding (from 24 to 60 h) and pH adjustment (from 24 to 72 h) resulted in a high NFBC yield (7.6 g/L; **Fig. 1g**) due to promoted cell growth (**Fig. 1h**). Under this condition (condition 3), total NFBC yield was double than that under condition 2 (**Fig. 1d**), but the specific NFBC production rate was only slightly increased (0.47 g/g_cdw_ h at 36 h; **Fig. 1h**). The feeding method (7 mL/h; 250 mL of 6.25% [w/v] glucose and 6.25% [w/v] fructose solution) under condition 3 was determined empirically to ensure that the sugars were almost consumed after 72 h (**Fig. 1i**). Other related data are shown in Fig. S2g-i.

The culture results of varying sugar concentration conditions are shown in Fig. S3. When the feeding amount and rate were increased (500 mL at 14.0 mL/h, condition S1), GA was not fully consumed (**Fig. S3a**), leading to a large pH drop (**Fig. S3b**), slower growth (**Fig. S3c**) and low NFBC yield (5.3 g/L; **Fig. S3a**). A large amount of fructose remained at 72h (**Fig. S3d**). When the initial sugar concentration was doubled (2.5% [w/v] glucose and 2.5% [w/v] fructose, condition S2), GA accumulation within the initial 24 h was doubled (**Fig. S3e**), leading to a significant decrease of pH (**Fig. S3f**) and cell growth (**Fig. S3g**). More than half of the fructose remained in the media at 72h (**Fig. S3h**), and the final NFBC yield (3.0 g/L; **Fig. S3e**) was lower than that under condition 2 (3.5 g/L; **Fig. 1d**).

Based on the above results, an initial sugar concentration of 2.5% (w/v) (1.25% [w/v] glucose and 1.25% [w/v] fructose) and constant feeding of 12.5% (w/v) sugar solution (6.25% [w/v] glucose and 6.25% [w/v] fructose) at 7.0 mL/h were used as the basic fed-batch culture method for subsequent experiments.

### NFBC production by the *gcd1* knockout mutant

As indicated above, rapid synthesis of GA decreased the pH, thereby inhibiting bacterial growth. This reaction reduced the final NFBC yield due to glucose depletion. To further investigate the impacts of GA synthesis and assimilation on bacterial growth and cellulose production, we constructed a *gcd* knockout mutant using the markerless gene deletion method (Kostner 2017; Bimmer 2022). To improve the efficiency of recombinant colony screening, we incorporated a blue–white selection system into our gene deletion method by introducing indigoidine synthesis-related genes (*bpsA* and *pptA*), as previously described (Rezuchova et al. 2018).

During the screening of the first recombination, a dark blue colony was selected from an agar plate containing a low concentration of Amp (100 μg/mL) to minimize the risk of off-target plasmid insertion into the genome. During this process, many background colonies (non-recombinant strains) were formed on the agar plate, and the colony in which the first recombination had occurred was selected based on its color (**Fig. 2a**, left). Only one blue colony was obtained from two agar plates inoculated with 150 μL of cell suspension after electroporation, indicating that the efficiency of first recombination was very low. Recombination efficiency could not be calculated because of the large number of background colonies. The target colony was inoculated again on an agar plate containing Amp (100 μg/mL) to separate it from the background colonies (**Fig. 2a**, center). The separated blue colony was subjected to second screening on an agar plate containing FC (240 µg/mL), and only white colonies were observed (**Fig. 2a**, right). The colony with the desired gene deletion was selected via PCR (**Fig. 2b** and c) and confirmed via Sanger sequencing. To the best of our knowledge, this study is the first to successfully use the indigoidine-based blue–white selection method for *Komagataeibacter* species.

**Fig. 2.**
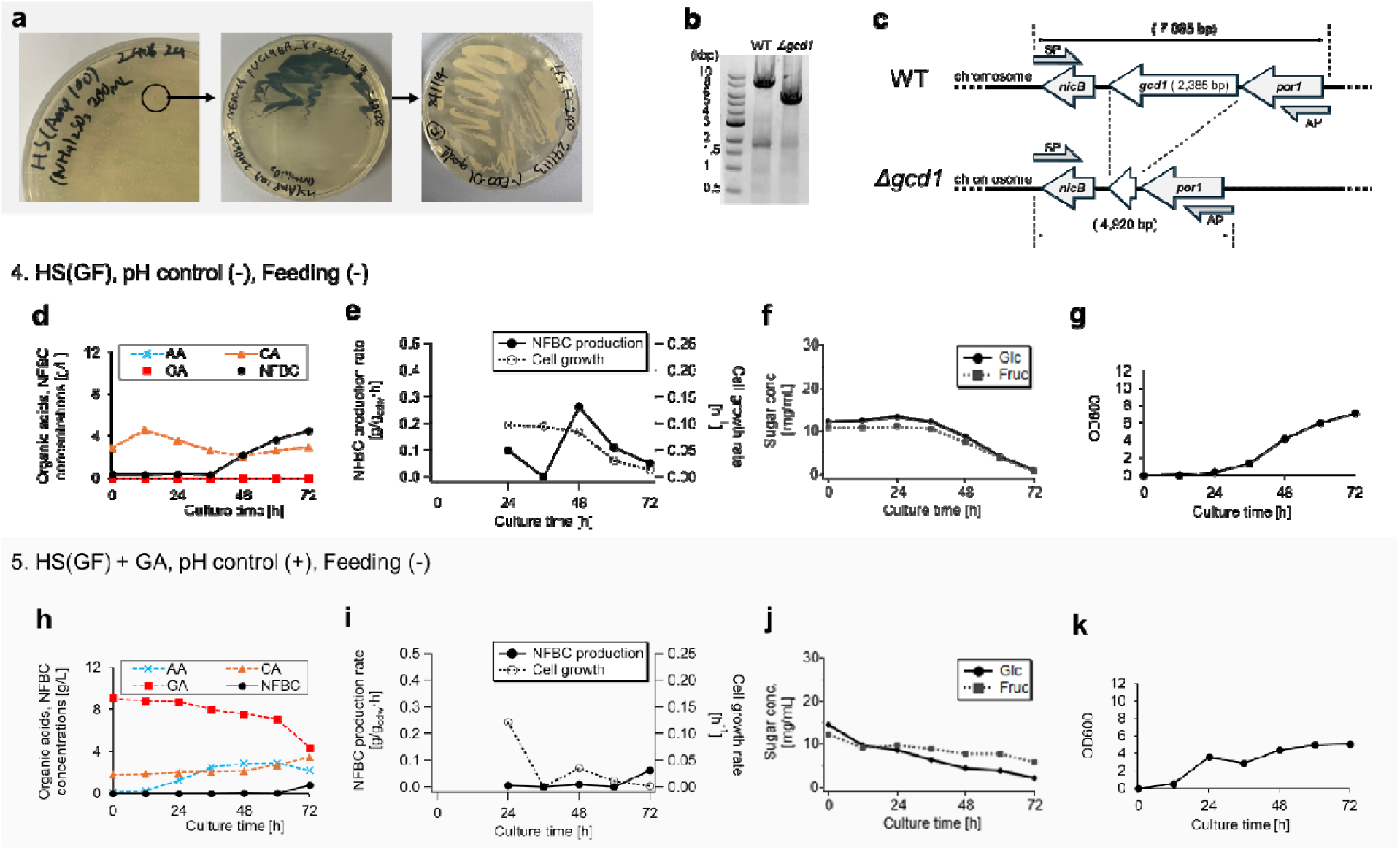
Preparation and culture of the *K*. *intermedius* glucose dehydrogenase (*gcd*)-deficient (Δ*gcd1*) strain. (**a**) Screening of target recombinant colonies via blue–white selection using indigoidine production as an indicator. (**b**) Agarose gel electrophoresis of the polymerase chain reaction (PCR) products amplified in the chromosomal region including *gcd1*. (**c**) Target chromosomal region for PCR before (wild-type [WT], top) and after (Δ*gcd1*, bottom) gene deletion via two successive homologous recombinations. SP, sense primer named SP_gcd1_upper_template; AP, antisense primer named AP_gcd1_lower_template (**Table S2**). (**d** and **h**) Time courses of NFBC and organic acid accumulation in the culture media (extracellular). (**e** and **i**) Specific NFBC production and growth rates estimated from the OD_600_ values. Data points at 12 h were omitted due to low OD values. (**f** and **j**) Glucose and fructose concentrations in the culture media. Culture conditions were as follows: pH regulation (24–72 h) using 2 M NaOHaq and 0.5 M H_2_SO_4_ (condition 4; **d**–**g**); 10 g/L GA was initially added to the HS(GF) medium and pH was controlled at 24–72 h (condition 5; **h**–**k**). All cultures involved the Δ*gcd1* strain

When the *gcd*-deficient strain (Δ*gcd1*) was cultured in the HS(GF) medium without pH control, no GA accumulation or decrease in pH was observed (**Figs. 2d** and **S4a**). Total NFBC yield (4.53 g/L; **Fig. 2d**) was significantly increased (by 85% relative to condition 1 [2.45 g/L; **Fig. 1a**] and 28% relative to condition 2 [3.53 g/L; **Fig. 1d**]) compared to that in the wild-type cultures. However, NFBC production rate (maximum: 0.26 g/g_cdw_ h; **Fig. 2e**) was lower than that of the wild-type (maximum: 0.28 g/ g_cdw_ h under condition 1 [**Fig. 1b**] and 0.44 g/g_cdw_ h under condition 2 [**Fig. 1e**]). Unlike in the wild-type strain, glucose and fructose were simultaneously consumed in the latter half of the culture period (**Fig. 2f**). The specific cell growth rates at 36–48 h were remarkably increased, and final OD_600_ (7.11; **Fig. 2g**) was increased by 47% compared to that under condition 2 in the wild-type strain (4.85; **Fig. S2d**), whereas the start of cell growth was delayed in Δ*gcd1* (**Fig. 2g**) compared to that in the wild-type culture (**Fig. 1a**, **d**, and **g**).

To investigate the effects of GA metabolism on cell growth and cellulose synthesis, we examined the behavior of the Δ*gcd1* strain in a medium containing glucose, fructose, and GA (10 g/L sodium gluconate; condition 5; **Fig. 2h**). Unexpectedly, the specific NFBC production rate was markedly low (**Fig. 2i**), even though Δ*gcd1* started to take up sugars and grow at an earlier culture time (**Fig. 2j** and k), and final OD_600_ (5.07; **Fig. 2k**) was comparable to that in the wild-type (4.85 under condition 2; **Fig. S2d**). Notably, GA uptake by Δ*gcd1* was much slower than that by the wild-type, and acetic acid accumulation was observed in the culture medium (**Fig. 2h**). Other related data are shown in Fig. S4.

### Effects of Csl on *K. intermedius* culture and NFBC production

Csl is a fraction obtained from corn starch rich in organic nitrogen, amino acids (especially alanine), vitamins, and other nutrients. It is used in industrial fermentation processes instead of expensive nitrogen sources, such as peptones. In this study, we obtained a batch of Csl produced in Japan (Kougo starch, Chiba, Japan) and used it in various cultures. Organic and amino acid concentrations in Csl were measured (**Fig. S5** and **Table S3**). The organic acid component included a high concentration of lactic acid (70.7 g/L) and low concentration of acetic acid (10.2 g/L; **Table S3**).

In wild-type *K*. *intermedius* culture, lactic acid in the medium was rapidly consumed (**Fig. 3a**). GA accumulation was lower (**Fig. 3a**) and exhibited better cell growth than that in the HS(GF) culture (**Figs. 3b** and **S6a**). Both GA and lactic acid were rapidly consumed, resulting in a spontaneous increase in pH, even without pH control (**Fig. S6b**). Such pH stability and buffering capacity are possibly beneficial for bacterial growth, facilitating active sugar uptake and metabolic activities (**Figs. 3c** and **S6c**). Under pH uncontrolled, no feeding condition, final NFBC yield (4.99 g/L; **Fig. 3a**) was doubled compared to that in the HS(GF) medium (2.45 g/L) (**Fig. 1a**).

**Fig. 3.**
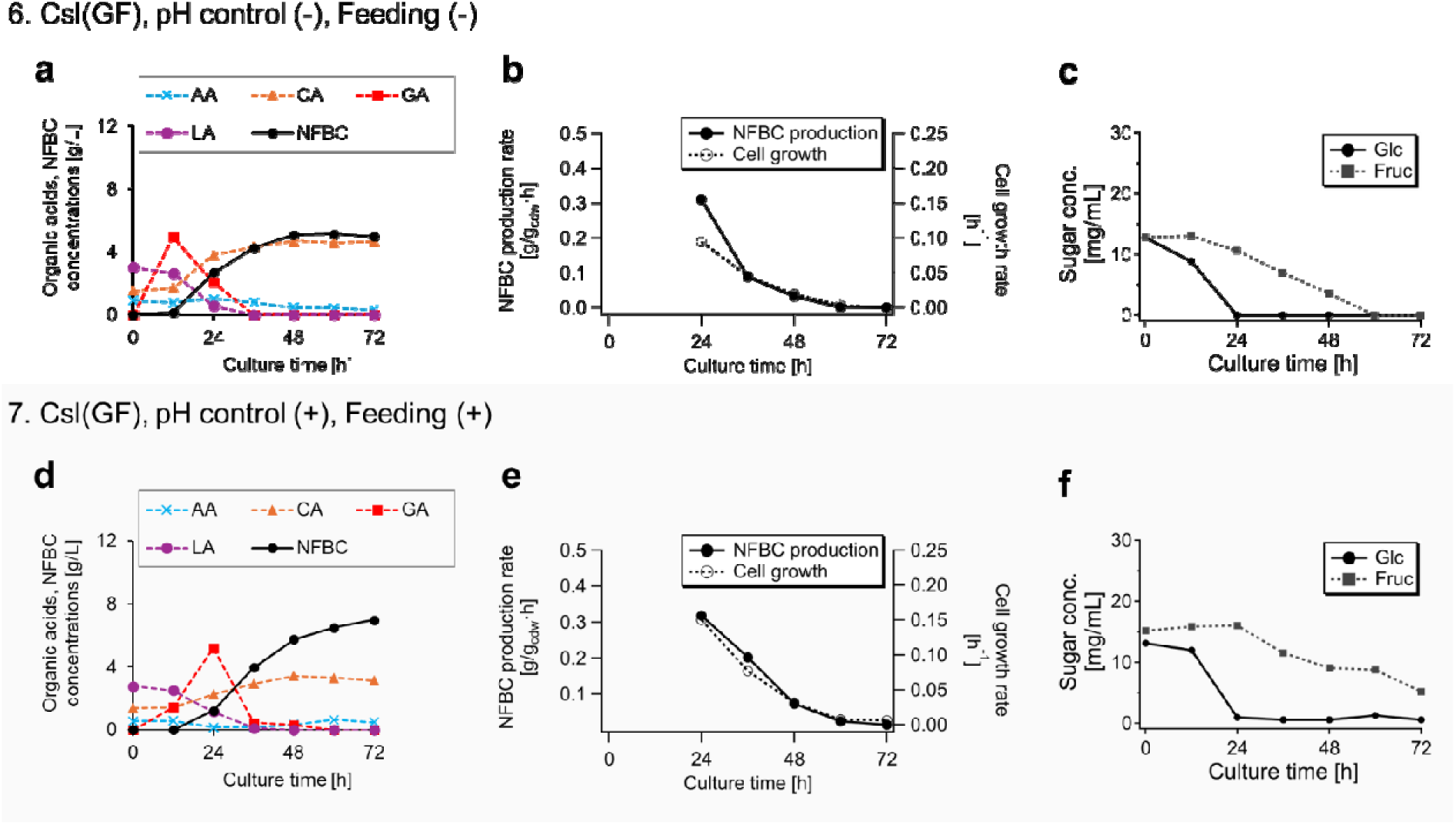
Effects of corn steep liquor (Csl) on *K*. *intermedius* culture. (**a** and **d**) Time courses of NFBC and organic acid accumulations in various culture media. (**b and e**) Specific NFBC production and growth rates estimated from the OD_600_ values. Plots at 12-h periods were omitted as the corresponding OD values were too low. (**c and f**) Sugar concentrations in the culture media. Culture conditions were as follows: Normal batch culture without pH control or feeding (condition 6; **a–c**); a combination of pH regulation (24–72 h) and feeding (24–60 h) at a rate of 7.0 mL/h (condition 7; **d**-**f**). All experiments used the wild-type *K. intermedius* NEDO-01 strain in the Csl(GF) medium, with an initial sugar concentration of 2.5% (w/v) (1.25% [w/v] glucose and 1.25% [w/v] fructose).

In the fed-batch condition, NFBC yield increased (7.0 g/L; **Fig. 3d**) due to the prolonged cellulose synthesis and cell growth phases in the presence of sugar (**Fig. 3e**), whereas the final yield was slightly lower than that in the HS(GF) medium (7.6 g/L; **Fig. 1g**). Although the maximum specific NFBC production rate was lower than that in the HS(GF) medium, a comparable NFBC yield was obtained (**Fig. 3d** and e), due to higher cell density (**Fig**. **S6d**). Fructose was not fully consumed throughout the culture period (**Fig. 3f**), and the respiration was more active after 48 h (**Fig. S6e** and **f**) compared to that in condition 6.

The presence of Csl also affected bacterial cell length. Microscope images and culture data in varying Csl or lactic acid concentrations are shown in Fig. S7.

### NFBC production from molasses and Csl

Use of natural resources, such as agro-industrial by-products, as culture medium components for BC production can reduce the environmental impact. Therefore, in this study, we performed a fed-batch culture using Csl-based media with three types of molasses from different sources and quantities as carbon sources and evaluated the resulting NFBC yields and characteristics.

We analyzed the organic and amino acid compositions of three molasses samples (**Fig. S5**; **Tables S3** and **S4**). Two samples were derived from sugar beets (molasses A and B), and one was derived from sugarcane (molasses SC). Molasses A and B differed in purity, with molasses A exhibiting higher sugar purity than molasses B. Both molasses B and SC were relatively crude, containing various organic and amino acids (**Tables S4** and **S4**).

Fig. 4 shows the results of the fed-batch cultures, in which different types of molasses were used as the carbon source instead of glucose and fructose. In Csl(H) medium, although the start of NFBC production was delayed, the final NFBC yields, cell growth, and sugar consumption rate were comparable to those obtained in Csl(GF) medium (**Figs. 4a-c** and **S8a** and **b**). In Csl(B) medium, NFBC yield was lower than that in Csl(A) medium (4.95 g/L; **Fig. 4d**). Also, the start of cell growth, NFBC production, and sugar uptake were slower than thos in Csl(A) medium (**Figs. 4d-f** and **S8c** and **d**). These delays were possibly due to the presence of impurities or specific amino acids (e.g., formic acid) in molasses B that inhibited bacterial growth. In contrast, in Csl(SC) medium, the final NFBC yield was comparable to that in Csl(A) and Csl(GF) (6.84 g/L; **Fig. 4g**), even though the start of NFBC production and sugar uptake were also delayed (**Fig. 4h** and i). The culture in Csl(SC) medium showed better cell growth (**Fig. S8e**), which probably enhanced the total NFBC yield. The pH drop in these three conditions were comparable to that of Csl(GF) (**Fig. S8b**, d, f). The culture experiment results are summarized in **Table 2**.

**Fig. 4.**
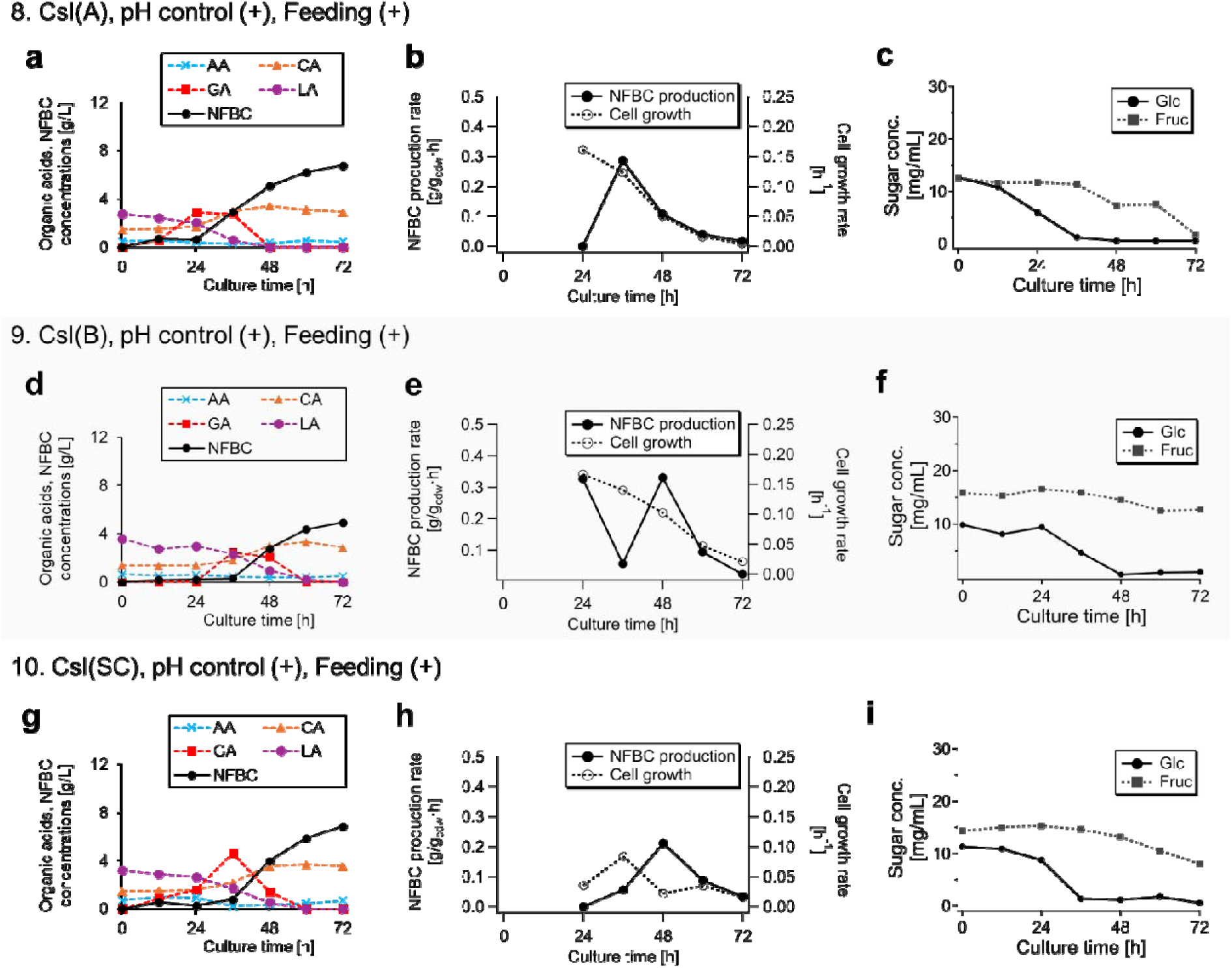
Fed-batch culture of *K*. *intermedius* in Csl-based media containing various molasses. (**a**, **d,** and **g**) Time courses of NFBC and organic acid accumulation in various culture media. (**b, e,** and **h**) Specific NFBC production and growth rates estimated from the OD_600_ values. Data points at 12 h were omitted due to low OD_600_ values. (**c, f,** and **i**) Sugar concentrations in the culture media. The following media were used under each condition: Csl(A) (condition 8; **a–c**), Csl(B) (condition 9; **d–f**), and Csl(SC) (condition 10; **g, h,** and **i**). All experiments were conducted using the wild-type *K. intermedius* NEDO-01 strain, with pH control (24–72 h) and feeding at a rate of 7.0 mL/h (24–60 h). Molasses concentrations in the culture media and feeding solutions were adjusted to achieve glucose concentrations of 12.5 and 62.5 g/L, respectively

**Table 2.** Summary of the results for NEDO-01 cultures using various methods and culture media.

| Condition | Method |  | Initial sugar<br>concentration<br>(%[w/v]) | Total sugar input |  | NFBC yield |  |  | Final OD<sub>600</sub> | Sugar consumption |  |
| --- | --- | --- | --- | --- | --- | --- | --- | --- | --- | --- | --- |
|  | pH<br>control | Feeding |  | Glucose<br>(g) | Fructose<br>(g) | vs. culture<br>volume (g/L) | vs. sugar<br>consumption<br>(%) | vs. total<br>sugar input<br>(%) |  | Glucose<br>(%) | Fructose<br>(%) |
| HS(GF), wild type |  |  |  |  |  |  |  |  |  |  |  |
| 1 | – | – | 2.5 | 62.5 | 62.5 | 2.45 | 9.8 | 9.8 | 4.92 | 100 | 100 |
| 2 | + | – | 2.5 | 62.5 | 62.5 | 3.53 | 14.4 | 14.1 | 4.85 | 100 | 95.2 |
| 3 | + | + | 2.5 | 93.8 | 93.8 | 7.59 | 23.1 | 21.2 | 7.64 | 100 | 83.9 |
| S1 | + | + | 2.5 | 125.0 | 125.0 | 5.32 | 15.8 | 11.8 | 7.05 | 95.0 | 54.7 |
| S2 | + | – | 5.0 | 125.0 | 125.0 | 3.00 | 8.3 | 6.0 | 3.78 | 96.5 | 46.7 |
| HS(GF), *Δgcd1* |  |  |  |  |  |  |  |  |  |  |  |
| 4 | – | – | 2.5 | 62.5 | 62.5 | 4.53 | 19.6 | 18.1 | 7.11 | 91.2 | 93.5 |
| 5 | + | – | 2.5 | 62.5 | 62.5 | 1.70 | 11.3 | 6.8 | 5.07 | 85.5 | 35.3 |
| Csl(GF), wild type |  |  |  |  |  |  |  |  |  |  |  |
| 6 | – | – | 2.5 | 62.5 | 62.5 | 4.99 | 20.0 | 20.0 | 8.22 | 100 | 100 |
| 7 | + | + | 2.5 | 93.8 | 93.8 | 6.98 | 22.6 | 19.5 | 9.81 | 100 | 72.4 |
| Csl(A), wild type |  |  |  |  |  |  |  |  |  |  |  |
| 8 | + | + | 2.1 | 93.8 | 67.2 | 6.78 | 22.1 | 23.5 | 9.61 | 100 | 86.0 |
| Csl(B), wild type |  |  |  |  |  |  |  |  |  |  |  |
| 9 | + | + | 2.4 | 83.9 | 80.4 | 4.95 | 27.5 | 15.6 | 8.57 | 93.5 | 18.2 |
| Csl(SC), wild type |  |  |  |  |  |  |  |  |  |  |  |
| 10 | + | + | 2.2 | 93.4 | 72.2 | 6.84 | 29.1 | 21.7 | 9.05 | 100 | 41.6 |

### Effects of various culture media on NFBC characteristics

Results of the structural analyses of NFBC samples are presented in Table 3. Native cellulose consists of extended molecular chains aligned parallel to form type-I cellulose crystals. These type I crystals are classified into two forms, I_α_ and I_β_, based on the differences in the relative positions of the molecular chains (Hebert and Muller 1974; Nishiyama et al. 2003; Nishiyama et al. 2002). Cellulose produced by *Komagataeibacter* species consists of a mixture of Iα and Iβ forms (Atalla and Vanderhart 1984). During molecular chain aggregation into cellulose ribbons, some regions remained non-crystalline, resulting in an amorphous cellulose structure. These amorphous regions affect the physicochemical properties of the resulting NFBC types, particularly their degree of crystallinity.

**Table 3.**
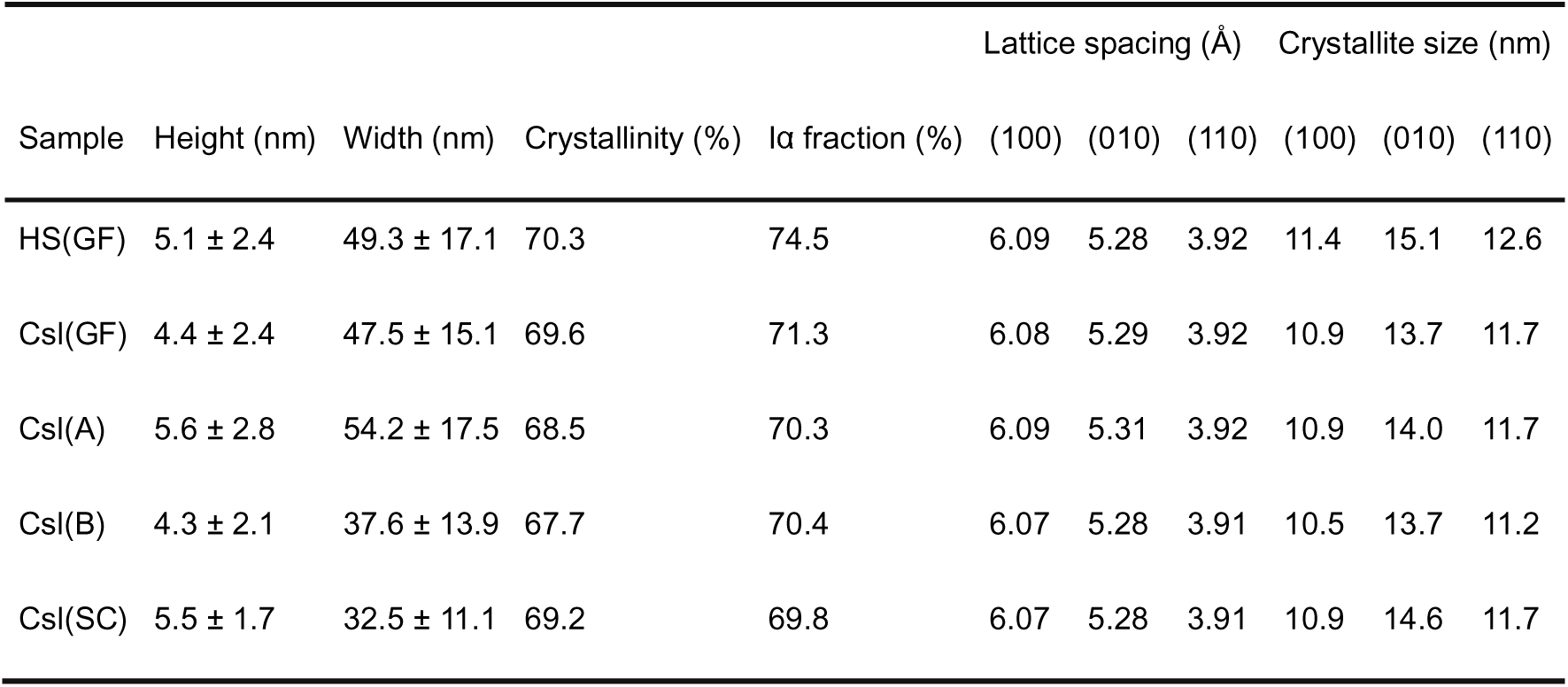
Results of the structural analysis of nanofibrillated bacterial cellulose (NFBC) types produced from various culture media (mean ± standard deviation [SD] values are presented for height and width).

| Sample | Height (nm) | Width (nm) | Crystallinity (%) | Iα fraction (%) | Lattice spacing ($\text{\AA}$) | | | Crystallite size (nm) | | |
| --- | --- | --- | --- | --- | --- | --- | --- | --- | --- | --- |
|  |  |  |  |  | (100) | (010) | (110) | (100) | (010) | (110) |
| HS(GF) | 5.1 ± 2.4 | 49.3 ± 17.1 | 70.3 | 74.5 | 6.09 | 5.28 | 3.92 | 11.4 | 15.1 | 12.6 |
| Csl(GF) | 4.4 ± 2.4 | 47.5 ± 15.1 | 69.6 | 71.3 | 6.08 | 5.29 | 3.92 | 10.9 | 13.7 | 11.7 |
| Csl(A) | 5.6 ± 2.8 | 54.2 ± 17.5 | 68.5 | 70.3 | 6.09 | 5.31 | 3.92 | 10.9 | 14.0 | 11.7 |
| Csl(B) | 4.3 ± 2.1 | 37.6 ± 13.9 | 67.7 | 70.4 | 6.07 | 5.28 | 3.91 | 10.5 | 13.7 | 11.2 |
| Csl(SC) | 5.5 ± 1.7 | 32.5 ± 11.1 | 69.2 | 69.8 | 6.07 | 5.28 | 3.91 | 10.9 | 14.6 | 11.7 |

Analysis of solid-state NMR spectra, measured using the ^13^C-CPMAS method, revealed that both the I_α_ fraction and degree of crystallinity were consistent across samples under different culture conditions. The I_α_ fraction was approximately 70%, with crystallinity of 70–75% (**Table 3**; **Fig. S9**). Additionally, its interplanar spacings calculated via WAXD analysis closely matched those of type I cellulose, with no observable change in crystal size across different cultivation conditions (**Table 3**; **Fig. S10**).

AFM analysis revealed that the characteristic ribbon-like morphology of NFBC was preserved under all culture conditions. The fibers exhibited flat ribbon-like structures with a mean height of 4–6 nm and width of 33–54 nm (**Table 3**; **Figs. 5** and **S11**). Notably, both height and width measurements exhibited substantial standard deviations, and no statistically significant differences were observed among the samples under different conditions.

**Fig. 5.**
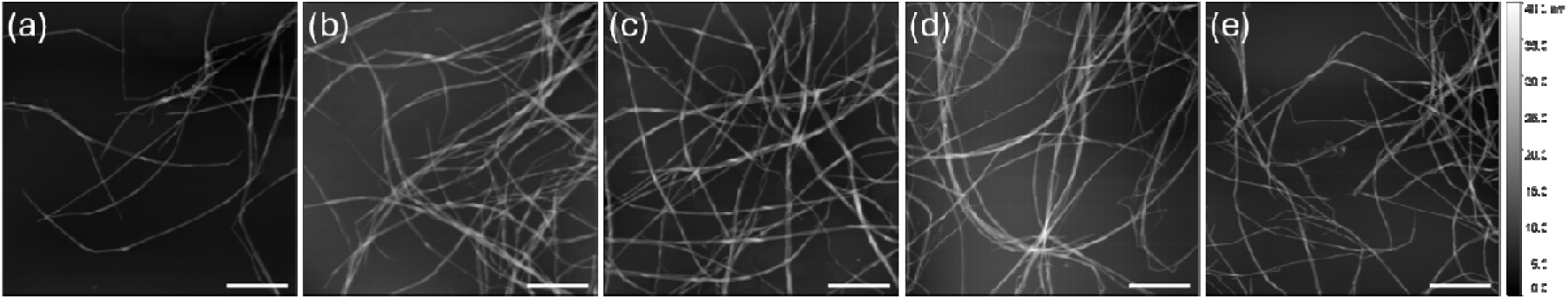
Atomic force microscopy images of the NFBC types prepared under different conditions. (a) HS(GF), (b) Csl(GF), (c) Csl(A), (d) Csl(B), and (e) Csl(SC). Scale bar in the image indicates 1 μm, and color bar on the right indicates height information

## Discussion

The conversion process of excess carbon into organic acid in microbial cultures is commonly known as overflow metabolism, a phenomenon well-characterized in *Escherichia coli* (Holms 1996). It relieves the metabolic pressure by diverting surplus carbon via alternative pathways. In contrast, oxidative fermentation involves the rapid oxidation of sugars or ethanol in the medium to organic acids via periplasmic dehydrogenases in acetic acid bacteria (Matsushita et al. 2004). Rapid conversion of glucose into GA is a key factor affecting BC production in cellulose-producing species, such as *Komagataeibacter* and *Novacetimonas* (Rezazadeh et al. 2020). Several studies have attempted to address this issue via *gcd* knockout experiments (Shigematsu et al. 2005; Kuo et al. 2015; Liu et al. 2020; Montenegro-Silva et al. 2024). However, only a few studies have investigated the associations among organic acid production, cellulose synthesis, and cell growth, particularly under varying culture conditions and in *gcd*-deficient strains. Therefore, in this study, we explored these interactions by analyzing carbon metabolism and organic acid synthesis under various culture conditions.

Wild-type NEDO-01 culture in the standard HS medium revealed that rapid GA accumulation via oxidative fermentation suppressed cell proliferation by decreasing the pH and depleting glucose, thereby reducing the NFBC yield (**Fig. 1**). NFBC production was enhanced by maintaining high cell proliferation via external pH control and appropriate carbon source feeding. However, because oxidative fermentation proceeds independently of cell growth, excessive glucose addition results in the overaccumulation of GA, which negatively affects the cell growth (**Fig. S3**). Therefore, for efficient NFBC production using the wild-type NEDO-01 strain, regulating the amount and timing of glucose feeding is crucial to prevent excessive GA accumulation and maintain conditions favorable for cell growth and cellulose synthesis.

In the Δ*gcd*1 strain culture, specific NFBC production rate was substantially lower than that in the wild-type culture, indicating a metabolic shift in which glucose-derived carbon was preferentially directed toward biomass formation rather than secondary metabolite (cellulose) synthesis. Nonetheless, total NFBC yield in the standard HS medium (condition 4) was higher in Δ*gcd*1, possibly due to enhanced biomass accumulation and prolonged cellulose production supported by sustained glucose availability (**Fig. 2d–g**). Notably, cell growth in Δ*gcd1* was delayed compared to that in the wild-type (**Fig. 2g** vs. **Fig. 1a**, **d**, and **g**), suggesting that early GA production and utilization contribute to cell growth initiation.

Indeed, under GA supplementation (condition 5), both glucose uptake and cell proliferation were initiated earlier (**Fig. 2h–k**), implying that GA functions as a trigger or enhancer of the central metabolic activity. The observed accumulation of acetic acid under this condition possibly reflects overflow metabolism, in which excess carbon is diverted from the central metabolic pathway and drained via pyruvate (Holms 1996). However, NFBC production was significantly reduced under this condition (**Fig. 2h** and **i**), indicating that carbon flux was redirected away from the cellulose biosynthesis pathway or that GA directly inhibited cellulose synthesis. In the wild-type culture, NFBC production coincided with a decline in GA accumulation during the early culture period (**Fig. 1a**, **d**, and **g**), further supporting that GA metabolism plays a regulatory role in cellulose synthesis initiation. However, further investigations are necessary to elucidate the mechanistic link between GA metabolism and cellulose biosynthesis.

Although several studies have explored the effects of *gcd* knockout on BC productivity (Shigematsu et al. 2005; Kuo et al. 2015; Liu et al. 2020; Montenegro-Silva et al. 2024), the results vary depending on the host strain and cultivation conditions. In static cultures, *gcd* knockout mutants of *K*. *xylinus* BPR2001 and *K*. *xylinus* BCRC1233 exhibit 1.4- and 1.7-fold increases in BC yield, respectively (Shigematsu et al. 2005; Kuo et al. 2015). Montenegro-Silva et al. reported a 5.77-fold increase in BC yield in a *gcd* knockout mutant of *K*. *sucrofermentans* ATCC700178 under optimized static culture conditions (Montenegro-Silva et al. 2024). In contrast, another study reported a 71.19% reduction in BC yield in a *gcd* knockout mutant of *K*. *xylinus* CGMCC 2955 under similar conditions (Liu et al. 2020). In our study, Δ*gcd1* mutant derived from *K*. *intermedius* NEDO-01 exhibited a 1.85-fold increase in NFBC yield in an STR without process optimization. Compared to the wild-type strain, Δ*gcd1* exhibited slower glucose consumption, suggesting that the intrinsic glucose uptake capacity of NEDO-01 is relatively low in the absence of periplasmic gcd. Future improvements in NFBC productivity can be achieved by upregulating glucose transport and enhancing BC biosynthesis via genetic regulation and optimization of the culture processes.

We also found that Csl reduced GA accumulation but activated the central metabolic pathway. Rapid consumption of lactic acid and concurrent accumulation of citric acid (**Fig. 3a** and **d**) indicated a metabolic shift in central carbon metabolism; lactic acid possibly served as a readily utilizable carbon and energy source, entering metabolism via pyruvate and subsequently feeding into the tricarboxylic acid cycle. Excretion (overflow) of citric acid indicated upregulated tricarboxylic acid cycle activity, supporting early phase cell growth. This metabolic boost reduced the need for glucose oxidation, thereby suppressing GA accumulation. Such a shift in metabolism possibly redirects the carbon flux toward biomass formation rather than cellulose biosynthesis, thereby decreasing the specific NFBC production rate.

Increased biomass in the Csl(GF) medium was possibly due to the increased individual cell size. After 72 h of culture in the Csl(GF) medium in baffled flasks, bacterial cells were significantly longer than those in the HS(GF) medium (**Fig. S7a** and **b**). A similar cell elongation effect was observed when ≥0.5% (w/v) lactic acid was added to the HS(GF) medium (**Fig. S7c**). Notably, NFBC yield decreased with increasing lactic acid concentration (**Fig. S7d**). These findings suggest that lactic acid plays a critical role in modulating central metabolism, potentially contributing to enhanced biomass production.

In fed-batch cultures using each molasses type combined with Csl, GA accumulation was similar to or slightly lower than that in the Csl(GF) medium, and pH fluctuations were smaller (**Fig. S8b**, **d** and **f**). Furthermore, fructose consumption was decreased in media containing molasses B and SC, resulting in high NFBC yields relative to the total sugar consumption (27.5% in Csl(B) and 29.1% in Csl(SC); **Table 2**). These results suggest that the amino and organic acids in molasses are preferentially utilized as carbon sources instead of fructose.

Analysis of solid-state NMR spectra, measured using the ^13^C-CPMAS method, revealed that both the I_α_ fraction and degree of crystallinity remained consistent across samples under different culture conditions. These values were also consistent with previous reports (Yamamoto and Horii 1994; Brouwer and Mikolajewski 2023), indicating that the NFBC types produced by *K. intermedius* NEDO-01 were structurally robust to metabolic variations induced by changes in the medium composition and culture method.

AFM observations were consistent with earlier reports that the presence of dispersing agents inhibits BC nanofiber bundling, promoting the formation of individual NFBC types (Kose et al. 2013; Tajima et al. 2017). Notably, NFBC fiber width varied depending on the cultivation conditions. This suggests that non-scaling properties, such as crystallinity, remain unaffected, whereas scaling properties, such as fiber dimensions, are sensitive to the culture conditions.

Width of the cellulose fibers produced by acetic acid bacteria correlates with the bacterial cell length (Yamanaka et al. 2000) due to the linear arrangement of cellulose synthase complexes along the cell surface (Brown et al. 1976). Accordingly, changes in cultivation conditions possibly influence bacterial metabolism, altering the cell length and NFBC fiber morphology. These findings highlight the potential to control the NFBC fiber width by regulating the bacterial cell length using customized culture strategies, which show great promise for the production of different NFBC types with tailored fiber dimensions.

## Conclusions

In this study, we quantitatively clarified the effects of GA production via oxidative fermentation, which is commonly observed in the *Acetobacteraceae* family, on cellulose production by *K. intermedius* NEDO-01 under different culture conditions in STRs. Notably, GA rapidly accumulated in the culture medium in proportion to the initial glucose concentration independent of bacterial growth, leading to a sharp drop in pH and glucose depletion. These effects primarily inhibited cell growth and reduced the cellulose yield. However, pH control and controlled feeding in STRs effectively mitigated these effects, resulting in a 3-fold increase in NFBC yield.

Interestingly, GA promoted the initiation of cell growth and redirected the glucose flux toward central carbon metabolism, while potentially inhibiting the cellulose synthesis pathway.

We assessed the potential use of agro-industrial by-products for NFBC production. These materials, particularly CSL, contain high levels of organic compounds and amino acids. Lactic acid in Csl was actively consumed by NEDO-01 and enhanced the cell growth. Although the use of such nutrient-rich by-products suppressed GA production and increased the cell growth rate, it reduced the specific NFBC production rate.

In Δ*gcd* strain cultures supplemented with GA, glucose uptake was as slow as fructose uptake, but carbon overflow was observed. This indicates that bypassing oxidative fermentation alone is insufficient to enhance cellulose production. Instead, enhancing glucose uptake and redirecting the glucose flux toward the cellulose biosynthesis pathway are critical to achieve high NFBC productivity.

Under optimized culture conditions, NFBC production was successfully enhanced using three types of molasses in combination with Csl. These cultures yielded relatively high NFBC concentrations, while maintaining consistent crystallinity and fiber width. Overall, our results suggest that NFBC of consistent quality can be reliably produced in STRs using *K. intermedius* NEDO-01 from various agricultural by-products.

## Data availability statement

The genome sequence of *Komagataeibacter intermedius* NEDO-01 has been deposited in GenBank under BioProject accession number PRJNA1303352 and BioSample accession number SAMN50505877. The genome data will be publicly released upon publication. Plasmids constructed in this study will be deposited to Addgene and made publicly available upon publication (Addgene ID: 246236). Also, datasets generated/analyzed in this study are available upon reasonable request from the corresponding author.

## Ethical approval

This study did not involve any human or animal experiments.

## Acknowledgments

Kota Noda, So Inoue, and Moemi Yoshimoto (Nitten): Organic acid analysis; Nozomi Takeda (GFC, Hokkaido University): Amino acid analysis; Naoya Nakagawa (Hokkaido University): NMR measurements; Eri Saito and Riku Furuse (Hokkaido University): Assistance with bioreactor experiments; Yuna Maeda (Hokkaido University): Assistance with gene knockout experiments; Risa Nakayama, Kahar Prihardi, and Chiaki Ogino (Kobe University): Technical support; Tokuo Matsushima and Ryo Serizawa (Kusano): Technical advice.

## Supplementary Material

### Supplementary Method

Method of plasmid construction (pUC19BAblue-Ki_gcd1)

## Supplementary Figure Captions

**Fig. S1** Plasmid design of pUC19BAblue-Ki_gcd1

**Fig. S2** Time course data () of cultures under conditions 1-3

**Fig. S3** Time course data of cultures under conditions S1-S2

**Fig. S4** Time course data of cultures under conditions 4-5

**Fig. S5** Balloon plot of organic acid and amino acid compositions in Csl and molasses

**Fig. S6** Time course data of cultures under conditions 6-7

**Fig. S7** Microscope images of bacteria cultured in varying Csl and lactic acid conditions

**Fig. S8** Time course data of cultures under conditions 8-9

**Fig. S9** Crystal structure analyses of NFBCs obtained under various culture conditions

**Fig. S10** Analyses of crystallinities for NFBCs obtained under various culture conditions

**Fig. S11** Histogram of fiber heights and width of NFBCs obtained under various culture conditions

## Supplementary Table Captions

**Table S1** Primers used in this study

**Table S2** Organic acid compositions of Csl, molasses A, B, and SC

**Table S3** Amino acid compositions of Csl, peptone, yeast extract, and molasses A, B, and SC

