## Supplemental figures and tables for "Modulation of gluconic acid metabolism enhances nanofibrillated bacterial cellulose production from agro-industrial by-products"

*Corresponding author:

Kenji Tajima

ORCID: 0000-0002-3238-813X

**Supplementary Method**

**Plasmid construction**

The plasmid for *gcd1* knockout, pUC19BAblue-Ki_gcd1, was constructed based on pUC19. The plasmids used in this study and the information (plasmid name, backbone, insert, source of insert, assembly method) are listed in Table S1.

Construction of pUC19BA

*E*. *coli* cytosine deaminase (*codA*, 1284 bp) and cytosine permease (*codB*, 1260 bp) genes (*codBA*, 2533 bp) for counter selection using fluorocytosine were amplified from the genomic DNA of *E*. *coli* BL21 (TaKaRa Bio Inc., Shiga, Japan) (GenBank accession number: CP010816.1). The vector fragment was amplified by PCR using a pUC19 fragment linearized with HindIII as the template. These two DNA fragments were assembled by InFusion method (TaKaRa Bio Inc.) according to the manufacturer’s instructions. The constructed plasmid was cloned using TaKaRa *E*. *coli* HST08 Premium Competent Cells (TaKaRa Bio Inc.).

Construction of pUC19BAblue-Ki_gcd1

Genomic DNA samples as PCR templates from the source organisms were extracted from the culture, using PrepMan Ultra Sample Preparation Reagent (ThermoFischer Scientific, Waltham, MA, USA). Indigoidine synthase gene (*bpsA*, 3849 bp, GenBank accession number: AB240063) was amplified by PCR from genomic DNA of *Streptomyces* *lavendulae* subsp. *lavendulae* NBRC 12340 (Purchased from Biological Resource Center [NBRC], National Institute of Technology and Evaluation [NITE], Tokyo, Japan). 4’-phosphopantetheinyl transferase gene (*pptA*, 738 bp) was amplified by PCR from genomic DNA of *Streptomyces* *mobaraensis* NBRC 13819 (Purchased from NBRC, NITE, Tokyo, Japan) (GenBank accession number: PRJDB448). Approximately 1200-bp region upstream and downstream of *gcd1* (upper flank and lower flank) were amplified by PCR, respectively. These DNA fragments were inserted into pUC19BA, linearized with PCR, by InFusion cloning according to the manufacturer’s instructions. The constructed plasmid was cloned using TaKaRa *E*. *coli* HST08 Premium Competent Cells.

**Supplementary Figures**

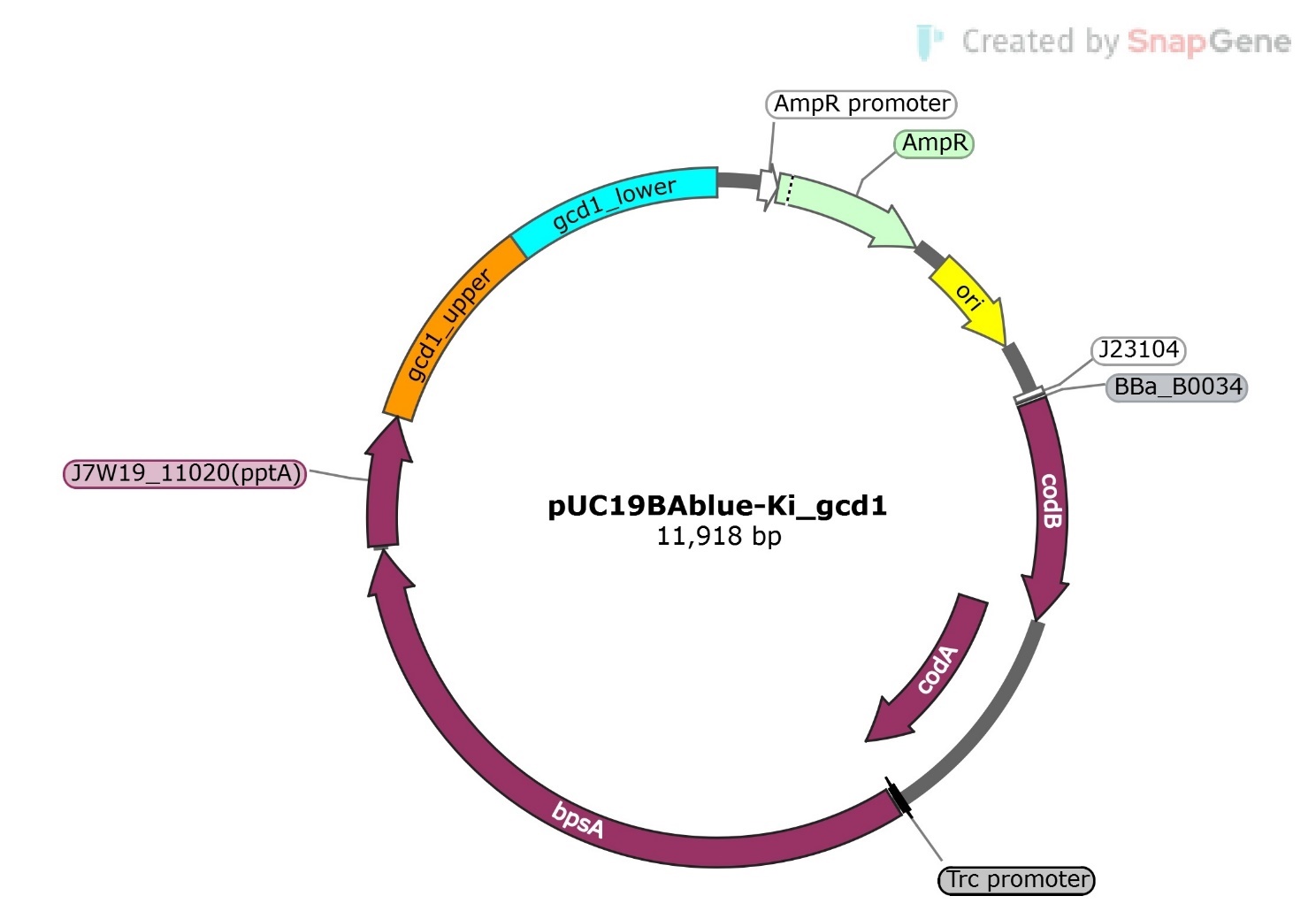

**Fig. S1** Plasmid design of pUC19BAblue-Ki_gcd1

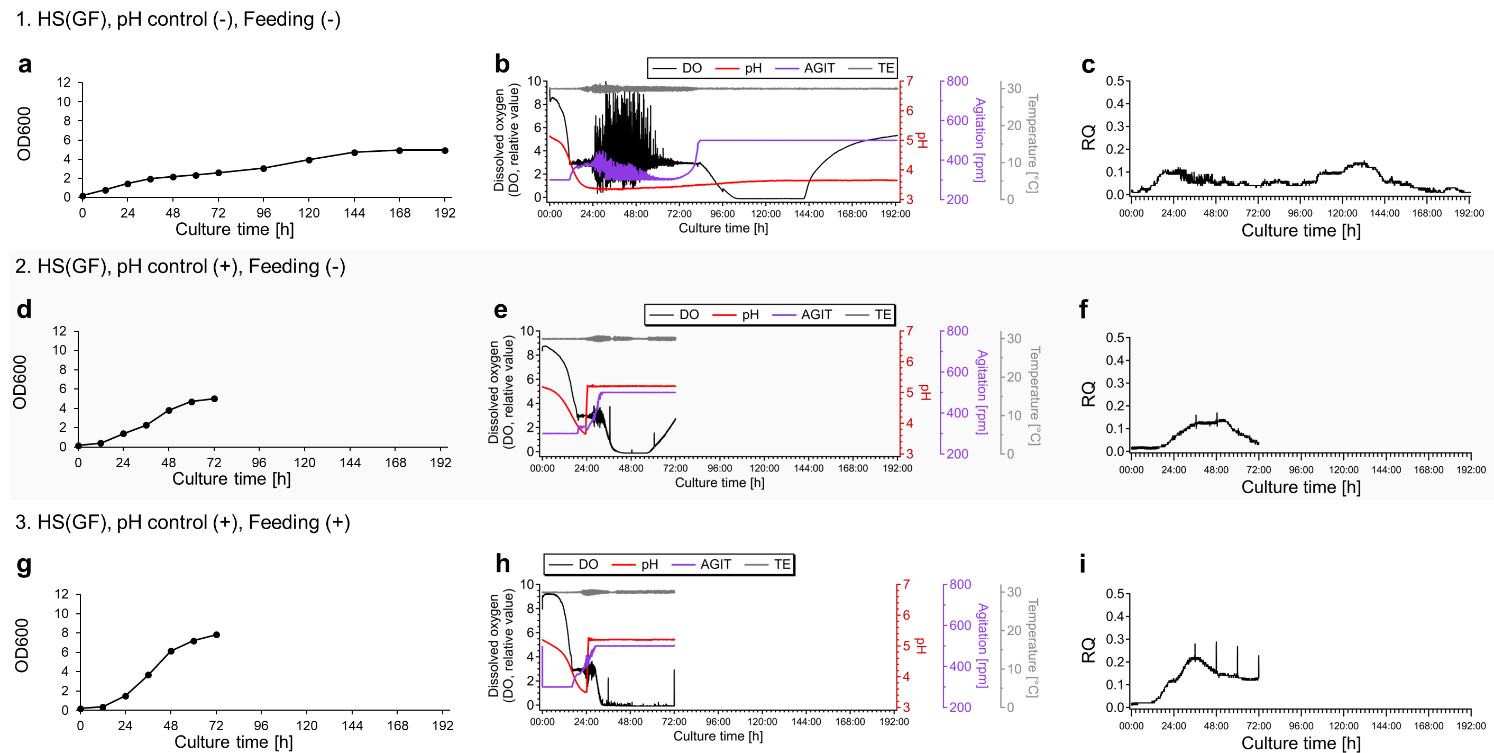

**Fig. S2** Time courses of optical density at 600 nm (OD_600_; **a**, **d**, and **g**), dissolved oxygen (DO), pH, agitation rate (AGIT), and temperature (TE) (**b**, **e**, and **h**), and respiratory quotient (RQ; **c**, **f**, and **i**) during cultivation. Culture conditions were as follows: Normal batch culture without pH control or feeding (condition 1; **a**–**c**); pH regulation (24–72 h) using 2 M NaOHaq and 0.5 M H_2_SO_4_ (condition 2; **d–f**); a combination of pH control (24–72 h) and feeding (24–60 h) at a rate of 7.0 mL/h (condition 3; **g–i**)

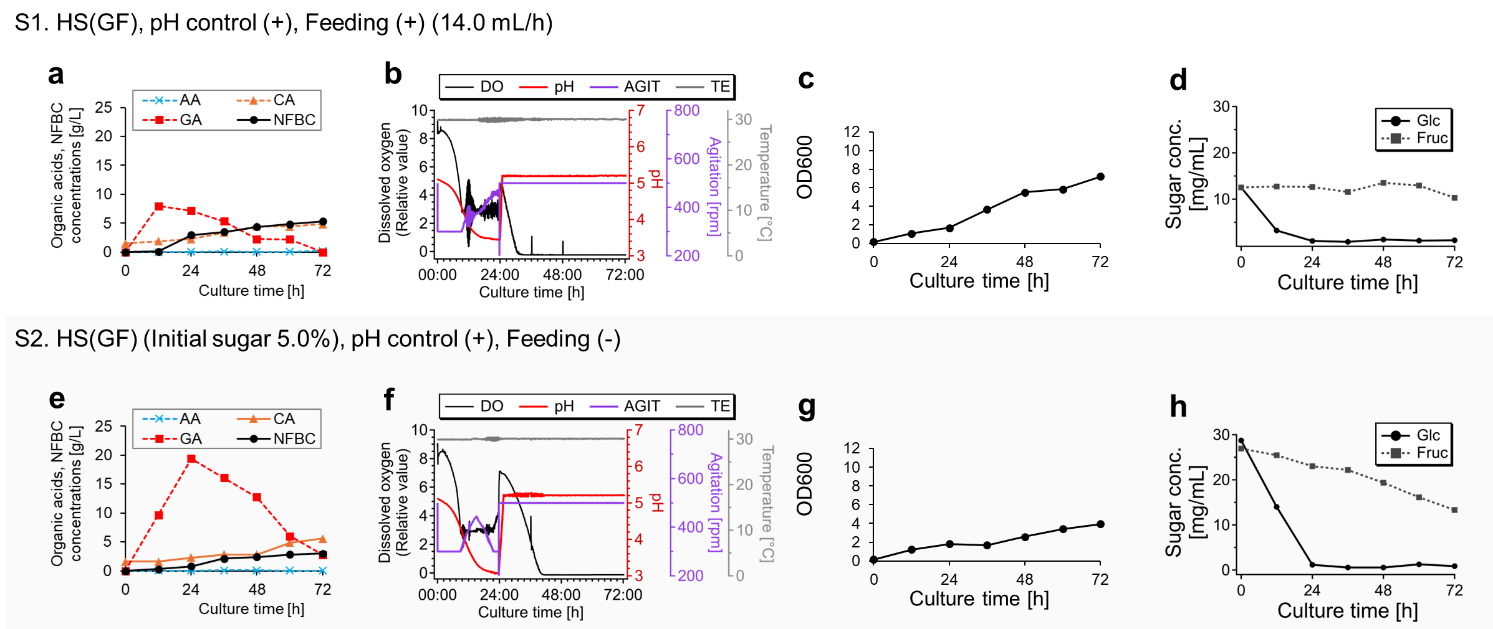

**Fig. S3** Time courses of NFBC and organic acid accumulation in the culture media (extracellular) (**a** and **e**), DO, pH, AGIT, and TE (**b** and **f**), OD600 (**c** and **g**), and glucose and fructose concentrations in the culture media (**d** and **h**). Culture conditions were as follows: a combination of pH regulation (24–72 h) and feeding (24–60 h) at a rate of 14.0 mL/h (condition S1; **a-d**); pH regulation (24–72 h) using 2 M NaOHaq and 0.5 M H_2_SO_4_, with the initial sugar concentration of 5% (w/v) (2.5% [w/v] glucose, 2.5% [w/v] fructose) (condition S2; **e–h**). All experiments used the wild-type *K.* *intermedius* NEDO-01 strain and Hestrin–Shramm (HS[GF]) medium

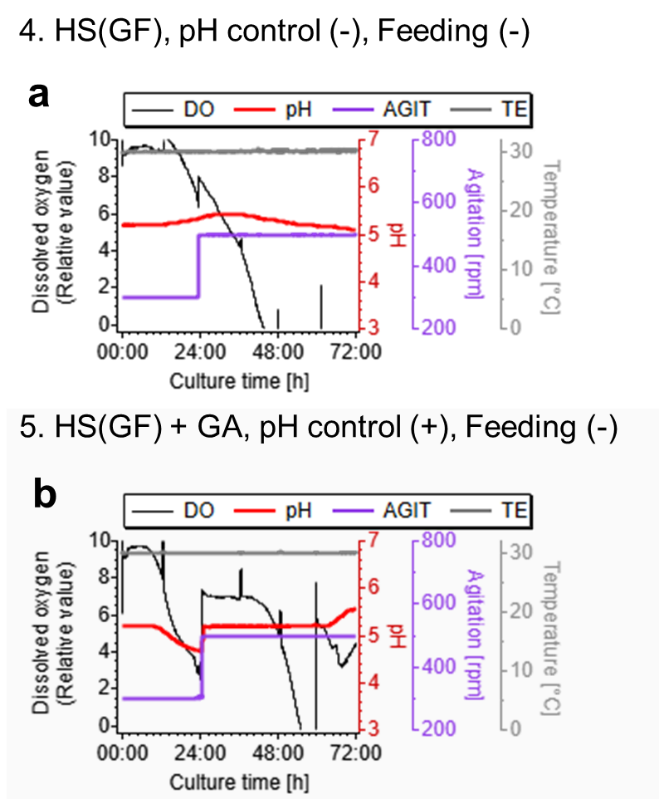

**Fig. S4** Time-courses of DO, pH, AGIT, and TE during cultivation. Culture conditions were as follows: pH regulation (24–72 h) using 2 M NaOHaq and 0.5 M H_2_SO_4_ (condition 4; **a**); 10 g/L GA was initially added to the HS(GF) medium and pH was controlled at 24–72 h (condition 5; **b**). All cultures used the *Δgcd1* strain and Hestrin–Shramm (HS[GF]) medium

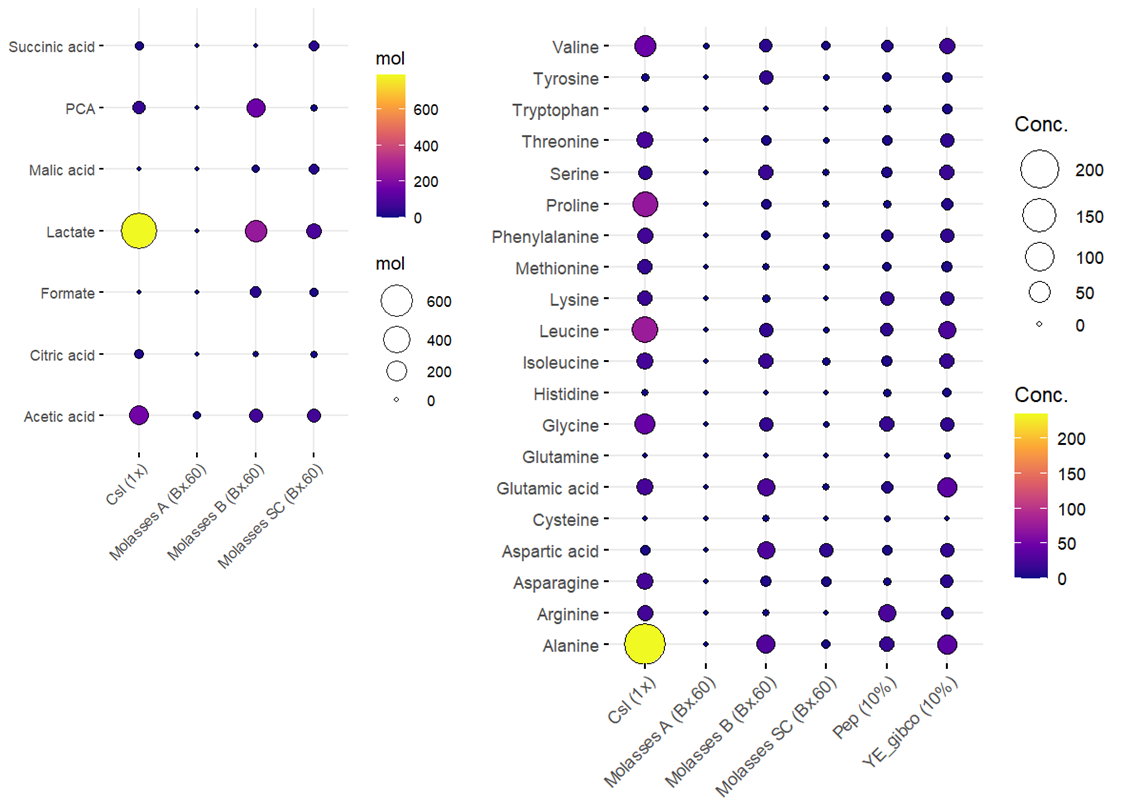

**Fig. S5** Organic acid and amino acid compositions in Csl and molasses

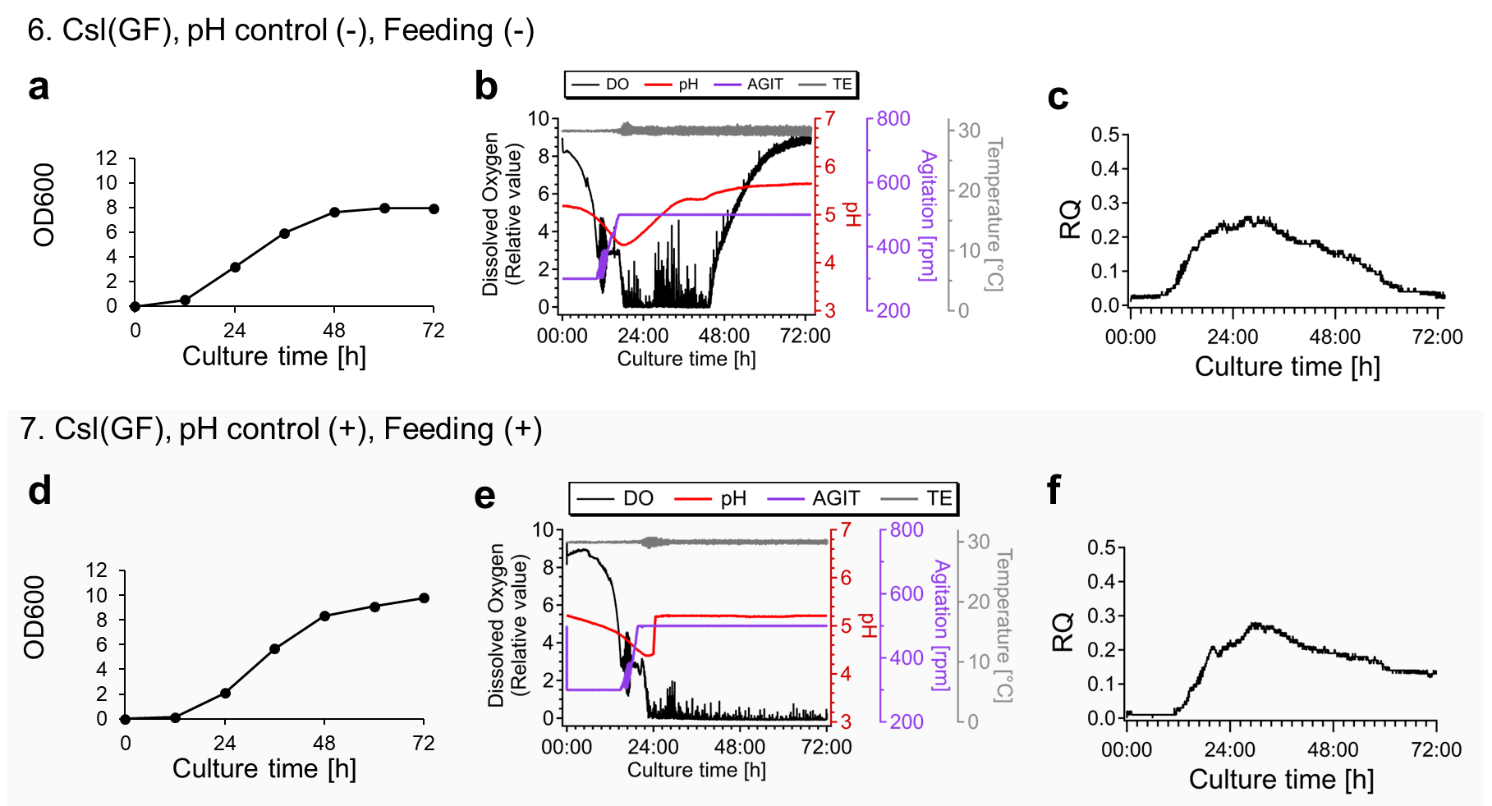

**Fig. S6** Time courses of OD_600_ (**a** and **d**), DO, pH, AGIT, and TE (**b** and **e**), and RQ (**c** and **f**) during cultivation. Culture conditions were as follows: Normal batch culture without pH control or feeding (condition 6; **a–c**); a combination of pH regulation (24–72 h) and feeding (24–60 h) at a rate of 7.0 mL/h (condition 7; **d**-**f**). All experiments used the wild-type *K. intermedius* NEDO-01 strain in the Csl(GF) medium, with an initial sugar concentration of 2.5% (w/v) (1.25% [w/v] glucose and 1.25% [w/v] fructose)

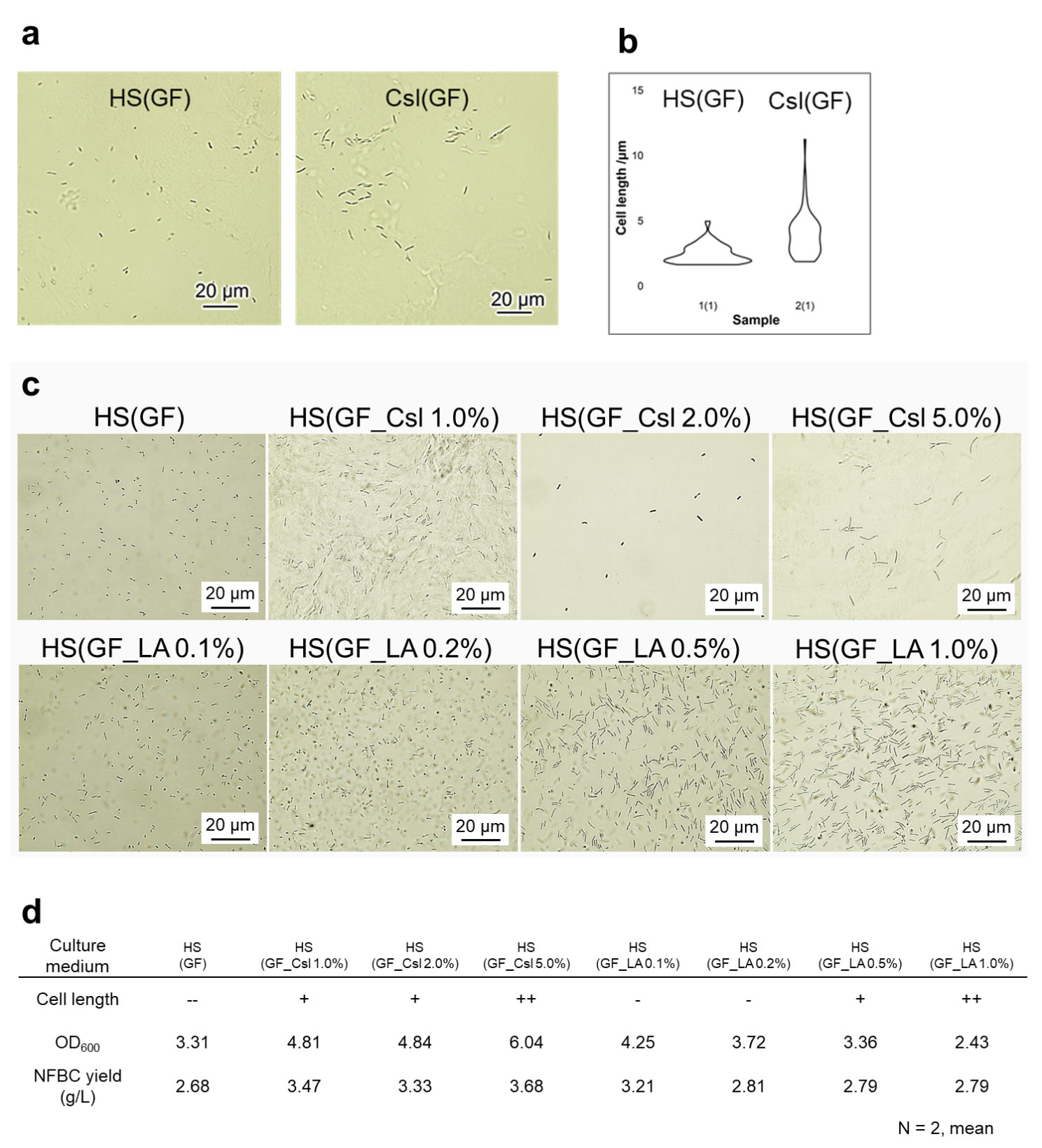

**Fig. S7** Microscope images of bacteria cultured in HS-based and Csl-based media (**a**), cell lengths distribution (**b**), microscope images of bacteria cultured in varying Csl or lactic acid concentrations based on HS media (**c**) and OD_600_ and NFBC yields in these conditions (**d**).

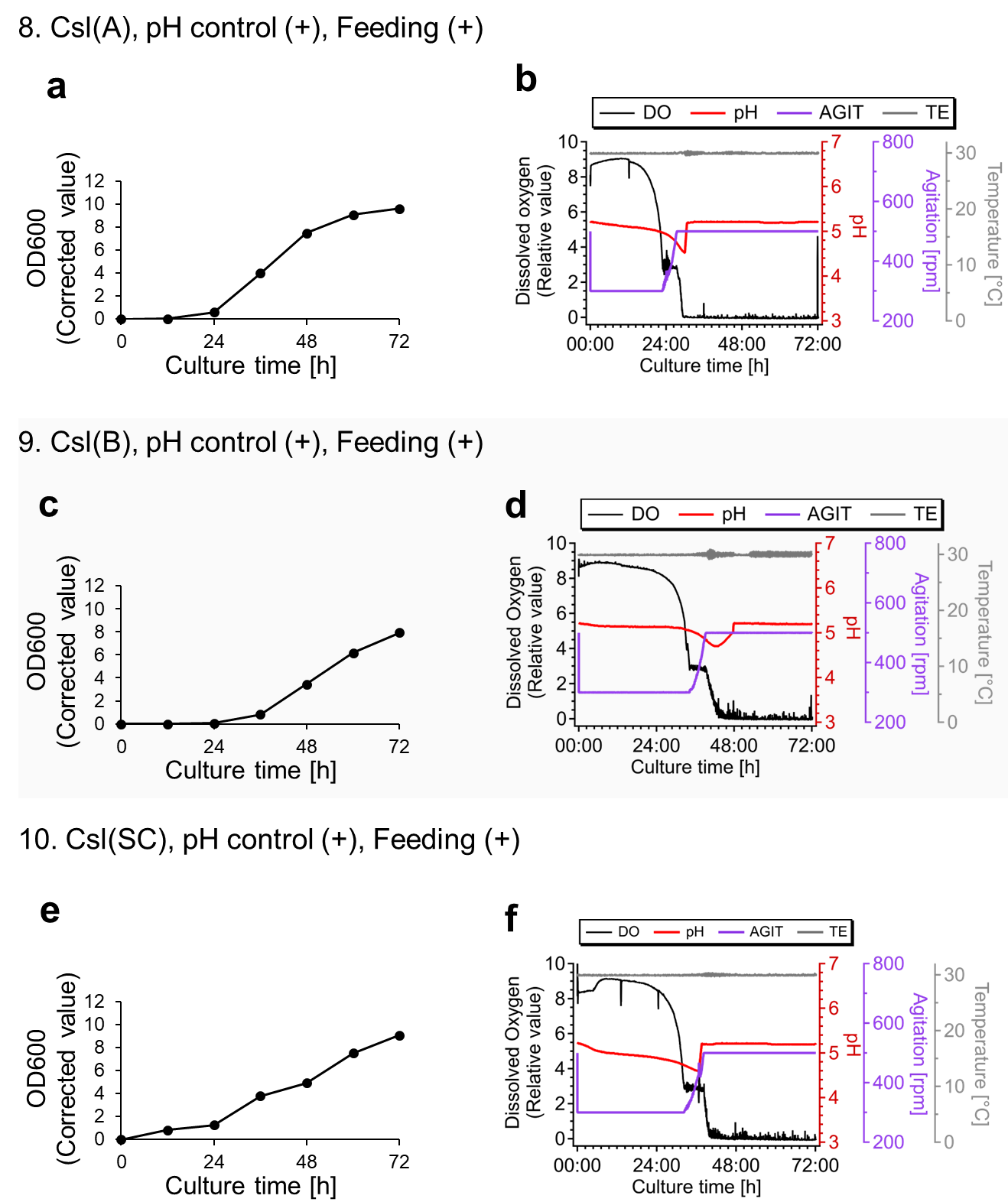
**Fig. S8** Time courses of OD_600_ (**a**, **c**, and **e**), and DO, pH, AGIT, and TE (**b**, **d**, and **f**) during Fed-batch cultivation. The following media were used under each condition: Csl(A) (condition 8; **a** and **b**), Csl(B) (condition 9; **c** and **d**), and Csl(SC) (condition 10; **e** and **f**). All experiments used the wild-type *K. intermedius* NEDO-01 strain, with pH control (24–72 h) and feeding at a rate of 7.0 mL/h (24–60 h). Molasses concentrations in the culture media and feeding solutions were adjusted to achieve glucose concentrations of 12.5 and 62.5 g/L, respectively

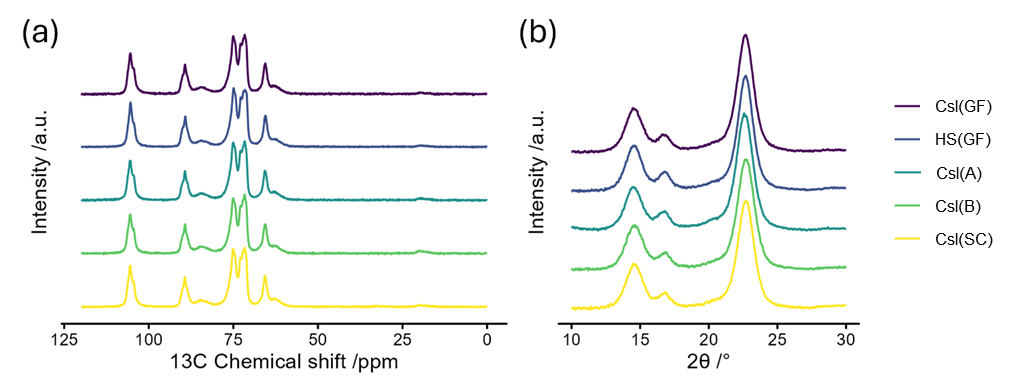

**Fig. S9** Crystal structure analyses of NFBCs obtained under various conditions: (**a**) solid-state CP-MAS ^13^C NMR spectra; (**b**) WAXD profiles

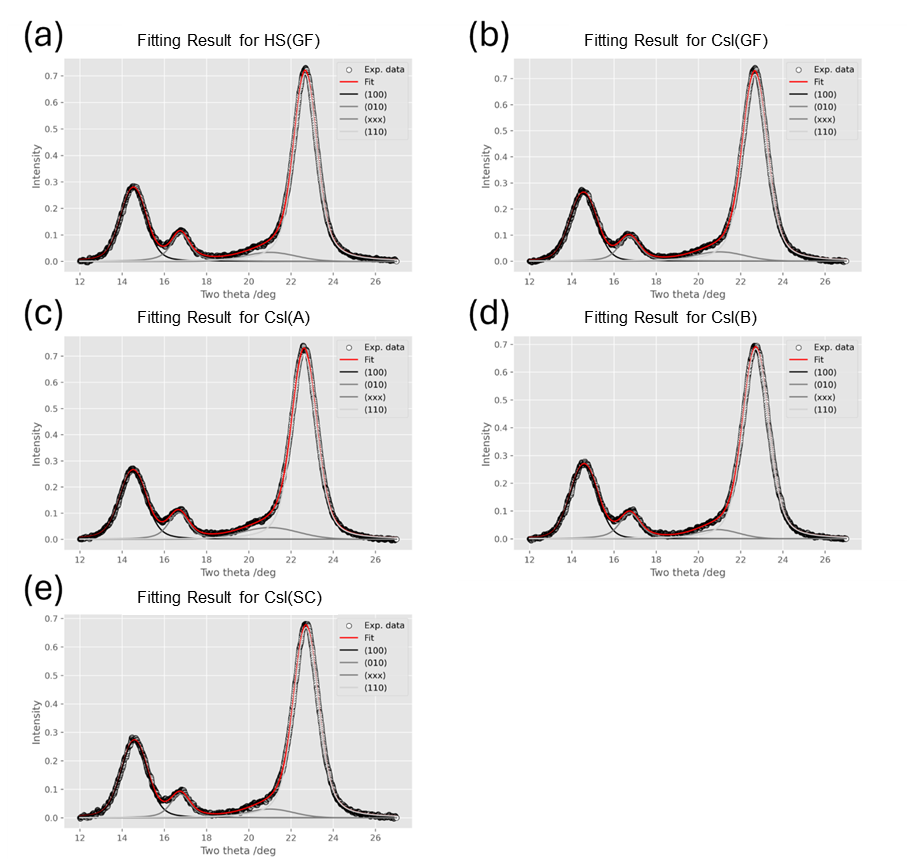

**Fig. S10** Analyses of crystallinities for NFBCs obtained under various conditions

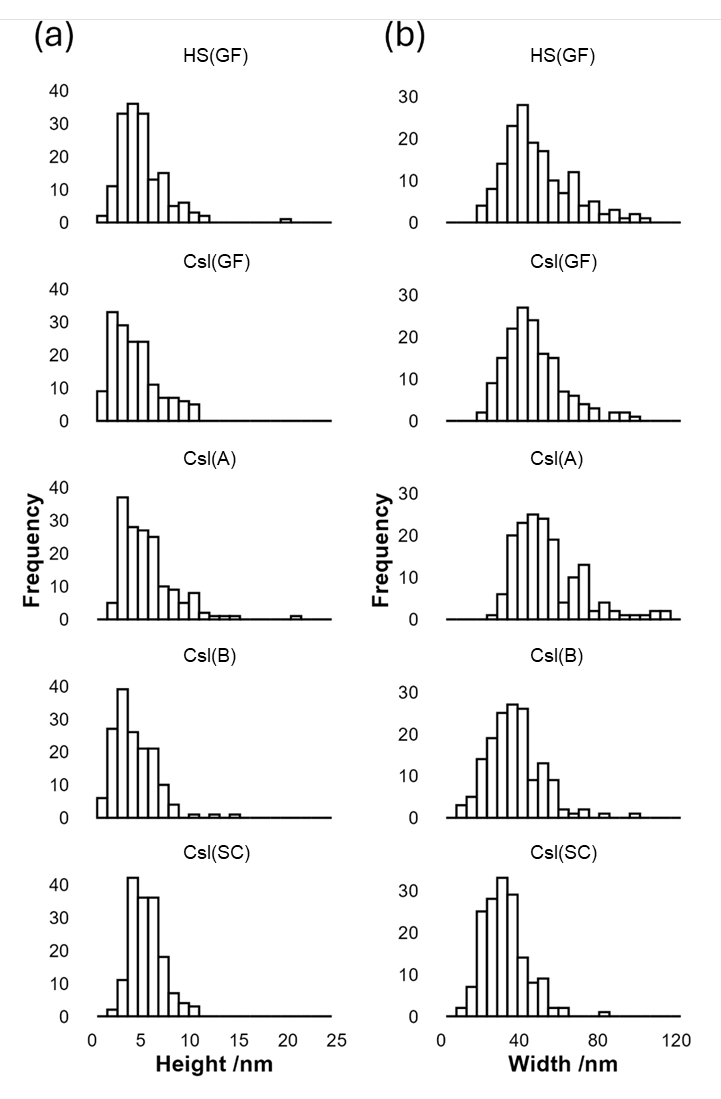

**Fig. S11** Histograms of fiber heights (**a**) and widths (**b**) of NFBCs obtained under various conditions

**Supplementary Tables**

**Table S1** Plasmids used in this study

| **Plasmid** | **Backbone** | **Insert** | **Source** | **Method** |
| --- | --- | --- | --- | --- |
| pUC19BA | pUC19 | *codBA* | *E*. *coli* BL21 (TaKaRa) (CP010816) | PCR +  InFusion cloning |
| pUC19BAblue-Ki_gcd1 | pUC19BA | *bpsA*  *pptA*  Upper flank region of *gcd1*  Lower flank region of *gcd1* | *Streptomyces lavendulae* subsp. *lavendulae* NBRC 12340 (*bpsA*) (AB240063)  *Streptomyces mobarensis* NBRC 13819 (*pptA*) (PRJDB448)  Komagataeibacter intermedius NEDO-01 (PRJNA1303352)  *K*. *intermedius* NEDO-01 | PCR +  InFusion cloning |

**Table S2** Primers

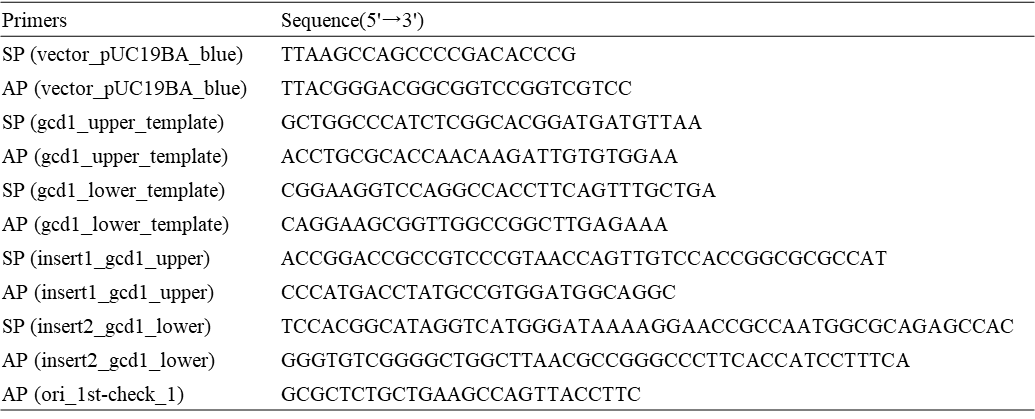

**Table S3** Organic acid compositions of Csl, Molasses A, B, and SC

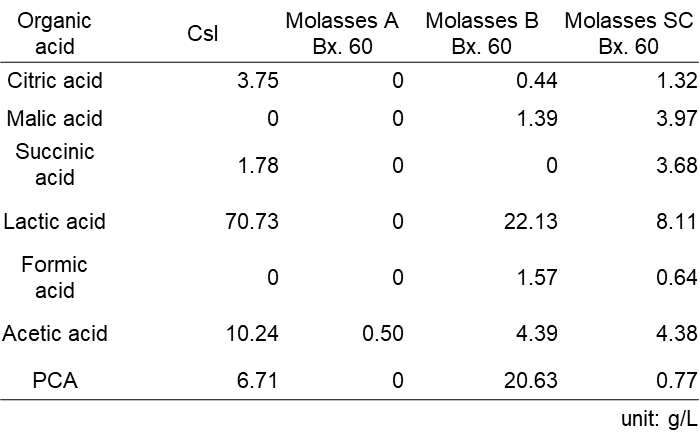

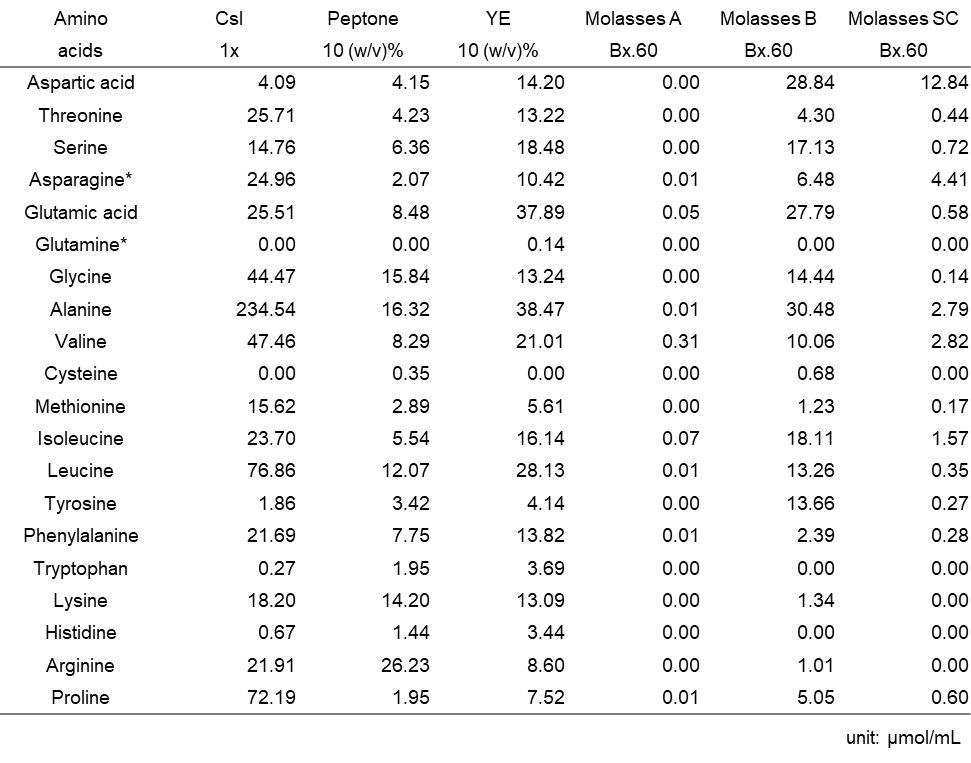
**Table S4** Amino acid compositions of Csl, peptone, yeast extract (YE), molasses A, B, and SC
